# Pyjacker identifies enhancer hijacking events in acute myeloid leukemia including *MNX1* activation via deletion 7q

**DOI:** 10.1101/2024.09.11.611224

**Authors:** Etienne Sollier, Anna Riedel, Umut H. Toprak, Justyna A. Wierzbinska, Dieter Weichenhan, Jan Philipp Schmid, Mariam Hakobyan, Aurore Touzart, Ekaterina Jahn, Binje Vick, Fiona Brown-Burke, Katherine Kelly, Simge Kelekci, Anastasija Pejkovska, Ashish Goyal, Marion Bähr, Kersten Breuer, Mei-Ju May Chen, Maria Llamazares-Prada, Mark Hartmann, Maximilian Schönung, Nádia Correia, Andreas Trumpp, Yomn Abdullah, Ursula Klingmüller, Sadaf S. Mughal, Benedikt Brors, Frank Westermann, Matthias Schlesner, Sebastian Vosberg, Tobias Herold, Philipp A. Greif, Dietmar Pfeifer, Michael Lübbert, Thomas Fischer, Florian H. Heidel, Claudia Gebhard, Wencke Walter, Torsten Haferlach, Ann-Kathrin Eisfeld, Krzysztof Mrózek, Deedra Nicolet, Lars Bullinger, Leonie Smeenk, Claudia Erpelinck, Roger Mulet-Lazaro, Ruud Delwel, Aurélie Ernst, Michael Scherer, Pavlo Lutsik, Irmela Jeremias, Konstanze Döhner, Hartmut Döhner, Daniel B. Lipka, Christoph Plass

**Affiliations:** Division of Cancer Epigenomics, German Cancer Research Center (DKFZ), Heidelberg, Germany; Division of Neuroblastoma Genomics, German Cancer Research Center (DKFZ), Heidelberg, Germany; Research Unit Apoptosis in Hematopoietic Stem Cells, Helmholtz Munich, German Research Center for Environmental Health, Munich, Germany; German Cancer Consortium (DKTK), partner site Munich, a partnership between DKFZ and University Hospital LMU Munich, Germany; Section of Translational Cancer Epigenomics, Division of Translational Medical Oncology, German Cancer Research Center (DKFZ); National Center for Tumor Diseases (NCT), NCT Heidelberg, a partnership between DKFZ and Heidelberg University Hospital, Heidelberg, Germany; Université de Paris Cité, Institut Necker Enfants-Malades (INEM), Institut National de la Santé et de la Recherche Médicale (Inserm) U1151, and Laboratory of Onco-Hematology, Assistance Publique-Hôpitaux de Paris, Hôpital Necker Enfants-Malades, Paris, France; Department of Internal Medicine III, University Hospital Ulm, Ulm, Germany; Heidelberg Institute for Stem Cell Technology and Experimental Medicine (HI-STEM gGmbH), Heidelberg, Germany; Division Systems Biology of Signal Transduction, German Cancer Research Center (DKFZ), Heidelberg, Germany; Division Applied Bioinformatics, German Cancer Research Center (DKFZ), Heidelberg, Germany; German Cancer Consortium (DKTK), Heidelberg, Germany; Medical Faculty and Faculty of Biosciences, Heidelberg University, Germany; Biomedical Informatics, Data Mining and Data Analytics, University of Augsburg, Augsburg, Germany; Department of Medicine III, University Hospital, LMU Munich, Munich, Germany; Department of Medicine I, Medical Center – University of Freiburg, Faculty of Medicine, Freiburg, Germany; Department of Hematology and Oncology, Medical Center, Otto-von-Guericke University, Magdeburg, Germany; Hematology, Hemostasis, Oncology and Stem Cell Transplantation, Hannover Medical School (MHH), Hannover, Germany; Leibniz Institute on Aging, Fritz-Lipmann-Institute, Jena, Germany; Interventional Immunology, Leibniz Institute for Immunotherapy, Regensburg, Germany; MLL Munich Leukemia Laboratory, Munich, Germany; Clara D. Bloomfield Center for Leukemia Outcomes Research, The Ohio State University Comprehensive Cancer Center, Columbus, OH, USA; Department of Hematology, Oncology, and Cancer Immunology, Charité, Universitätsmedizin Berlin, Berlin, Germany; Department of Hematology, Erasmus University Medical Center, Rotterdam, The Netherlands; Group Genome Instability in Tumors, German Cancer Research Center (DKFZ), Heidelberg, Germany; Department of Oncology, KU Leuven, Leuven, Belgium; Department of Pediatrics, Dr. von Hauner Children’s Hospital, University Hospital, LMU Munich, Munich, Germany; Faculty of Medicine, Otto-von-Guericke-University, Magdeburg, Germany

## Abstract

Acute myeloid leukemia with complex karyotype (ckAML) is characterized by high genomic complexity, including frequent *TP53* mutations and chromothripsis. We hypothesized that the numerous genomic rearrangements could reposition active enhancers near proto-oncogenes, leading to their aberrant expression. We developed pyjacker, a computational tool for the detection of enhancer hijacking events, and applied it to a cohort of 39 ckAML samples. Pyjacker identified *motor neuron and pancreas homeobox 1* (*MNX1*), a gene aberrantly expressed in 1.4% of AML patients, often as a result of del(7)(q22q36) associated with hijacking of a *CDK6* enhancer. *MNX1*-activated cases show significant co-occurrence with *BCOR* mutations and a gene signature shared with t(7;12)(q36;p13) pediatric AML. We demonstrated that *MNX1* is a dependency gene, as its knockdown in a xenograft model reduces leukemia cell fitness. In conclusion, enhancer hijacking is a frequent mechanism for oncogene activation in AML.

**Statement of significance:** This study examines the consequences of structural alterations and demonstrates that proto-oncogene activation by enhancer hijacking is an overlooked pathomechanism in AML. *MNX1* overexpression demonstrates that deletions on chromosome 7q can not only lead to haploinsufficiency, but also to activation of oncogenes by enhancer hijacking, providing a novel leukemogenic mechanism.

## Introduction

Acute myeloid leukemia (AML) is a disease characterized by a block in differentiation and uncontrolled proliferation of myeloid progenitor cells. AML is a very heterogeneous disease and has been divided into several subgroups based on recurrent cytogenetic alterations [e.g., t(15;17)(q24.1;21.2), inv(16)(p13.1q22) or t(8;21)(q22;q22.1)] and mutations (e.g., in *NPM1*, *TP53* or *CEBPA*) (1–3). Complex karyotype AML (ckAML) is a subtype with dismal prognosis and there is currently an incomplete understanding of the pathogenetic mechanisms driving this disease (4). ckAML is defined by the presence of at least three cytogenetic alterations, in the absence of any of the recurrent class-defining lesions. It accounts for 10-12% of all AML cases and is more frequent among older patients (4). ckAML samples often harbor *TP53* mutations, which are associated with a high frequency of chromothripsis, defined as the shattering of certain chromosomes and refusion in random order, resulting in highly rearranged chromosomes with loss of chromosomal material (5–7). Deletions in ckAML are more frequent than gains and the most common deletions affect chromosome arms 5q, 7q, 17p and 12p, while gains mostly occur on 8q, 11q and 21q (4,8,9). According to Knudson’s two-hit hypothesis, deletions in cancer usually lead to the complete inactivation of a tumor suppressor gene whose other copy is also inactivated, for example by a mutation. However, apart from *TP53* on 17p, the search for tumor suppressor genes with both copies inactivated in ckAML has been unsuccessful (4), and the current paradigm is that copy number alterations (CNAs) in ckAML lead to gene dosage effects driving tumorigenesis (10), where a higher (resp. lower) gene copy number results in a higher (resp. lower) gene expression.

Deletions of chromosomal segments on 7q are one of the most common structural alterations in AML (10%) (2,11). It is frequently seen in ckAML, but can also be found as a sole abnormality, where it is still associated with a poor prognosis (12). The clustering of these deletions in certain regions on 7q has been used for more than 20 years as an indication for the presence of a tumor suppressor gene within the minimally deleted region. However, the search for a gene with a second (epi)genetic hit has not been successful [reviewed by Inaba et al. (13)]. Consequently, the most plausible explanation for these highly recurrent clustered deletions is that they lead to haploinsufficiency of the genes in the deleted region, where the lower copy number results in reduced gene expression, and that this haploinsufficiency is sufficient to drive cancer. Of note, many haploinsufficient genes located in the deleted regions encode enzymes that regulate genome-wide epigenetic patterns or transcription factors such as *CUX1*, *EZH2*, *KMT2C* or *KMT2E* (13–15).

In addition to CNAs, structural variants (SVs) can create fusion proteins, or remove or create new enhancer-promoter interactions. For example, 5% of all AML cases harbor an inv(3)(q21q26.2) or a t(3;3)(q21;q26.2), which repositions the *GATA2* enhancer in close vicinity of *MECOM*, leading to aberrant *MECOM* expression and *GATA2* haploinsufficiency (16). A few other genes have been reported to be activated by enhancer hijacking in AML, including *BCL11B* in acute leukemias with a mixed phenotype (17) and *MNX1* in pediatric AML with t(7;12)(q36;p13) (18,19), but no systematic search for these events has been undertaken in AML to date. Since ckAML samples harbor many, often cytogenetically cryptic, genomic rearrangements, we hypothesized that some of them could lead to enhancer hijacking events, activating still-undiscovered oncogenes.

Recently, several computational methods have been developed to search for genes activated by enhancer hijacking. CESAM (20), SVExpress (21) and HYENA (22) perform a linear regression of gene expression depending on the presence of breakpoints nearby. These methods have successfully identified genes recurrently activated by enhancer hijacking, but they cannot detect genes activated in only a few samples. cis-X (23) can detect enhancer hijacking events in single samples using monoallelic expression, but this method is not very flexible and requires matched normal samples, which are rarely available for AML samples. NeoLoopFinder (24) follows a very different approach: it detects neo-loops in HiC data and does not use gene expression.

Here, we developed a new method, “pyjacker”, which detects putative enhancer hijacking events occurring in single samples, using RNA-seq and whole genome sequencing (WGS) without matched normal samples. We applied pyjacker to 39 ckAML samples using WGS and RNA-seq, and identified genes known to be activated by enhancer hijacking, as well as previously overlooked new candidate genes. We focused on *MNX1*, a gene encoding a homeobox transcription factor, which is mapped to chromosome band 7q36.3, that is located outside of the most commonly deleted regions found in AML with del(7q). We profiled 31 *MNX1*-expressing cases with WGS and discovered that del(7q) can lead to hijacking of the *CDK6* enhancer driving *MNX1* expression, resulting in a shared gene expression profile with pediatric AML with *MNX1* activation. We showed that *MNX1* knockdown reduces leukemic cell fitness in patient-derived xenograft (PDX) competition assays, demonstrating its essentiality.

## Results

### Pyjacker: detection of enhancer hijacking with WGS and RNA-seq

We developed pyjacker, a computational method to detect enhancer hijacking events occurring in single samples using WGS, RNA-seq and enhancer information, without the need for matched normal samples (Supplementary Table 1). For each gene, samples are divided into “candidate samples” which have breakpoints near the gene and “reference samples” which do not (see methods section for details). Reference samples are used to compute the mean and standard deviation of the expression of this gene in the absence of enhancer hijacking, and the candidate samples are tested for overexpression compared to this reference distribution (**Fig. 1A**). If a gene is activated by enhancer hijacking, we would expect only the rearranged allele to be expressed. Heterozygous SNPs are identified in the WGS data, and if these SNPs are covered in the RNA-seq data, pyjacker tests if the expression is monoallelic (**Fig. 1A**). Using the breakpoint information and a list of putative enhancers, pyjacker identifies enhancers coming close to the gene, and scores the event depending on the strength of the enhancers coming close to the gene. As enhancers are cell type-specific, we used in this study ChIP-seq data against H3K27ac and P300 from myeloid cell lines (see methods; Supplementary Table 2). This enhancer information can be omitted if it is not available. The overexpression, monoallelic expression and enhancer scores are combined into an empirical score which reflects how likely the gene is to be expressed because of a genomic rearrangement. The scores are aggregated across samples for each gene in order to give more weight to the recurrently activated genes. To estimate the false discovery rate (FDR), “null scores” are computed by only including the “reference samples”, and randomly assigning some of them to the “candidate samples”, thus reflecting the distribution of scores in the absence of enhancer hijacking. Finally, the Benjamini-Hochberg method is used to correct for multiple testing and provides a ranked list of genes putatively activated by a structural rearrangement, with corresponding FDR. Pyjacker is flexible and we provide an end-to-end nextflow pipeline to run pyjacker, starting from bam files. We note that fusion transcripts can also result in monoallelic overexpression, when the 3’ fusion partner is not normally expressed, although this would be a different mechanism than enhancer hijacking. Various methods can be used to detect fusion transcripts from RNA-seq data, like STAR-Fusion (25) or Arriba (26) and if a list of fusions is given as input to pyjacker, it will annotate candidate genes with the fusion status, allowing the identification of true enhancer hijacking events. Since pyjacker needs reference samples without breakpoints near a gene to estimate the reference expression distribution, it should be run with at least ten samples as input but works best with large cohorts. We tested pyjacker on ten AML cell lines, some of which have known enhancer hijacking events: *MECOM* in MOLM-1 and MUTZ-3 (16), *MNX1* in GDM-1 (27) and *MN1* in MUTZ-3 (28). Pyjacker correctly identified these four events with FDR<20% (Supplementary Table 3).

**Figure 1.**
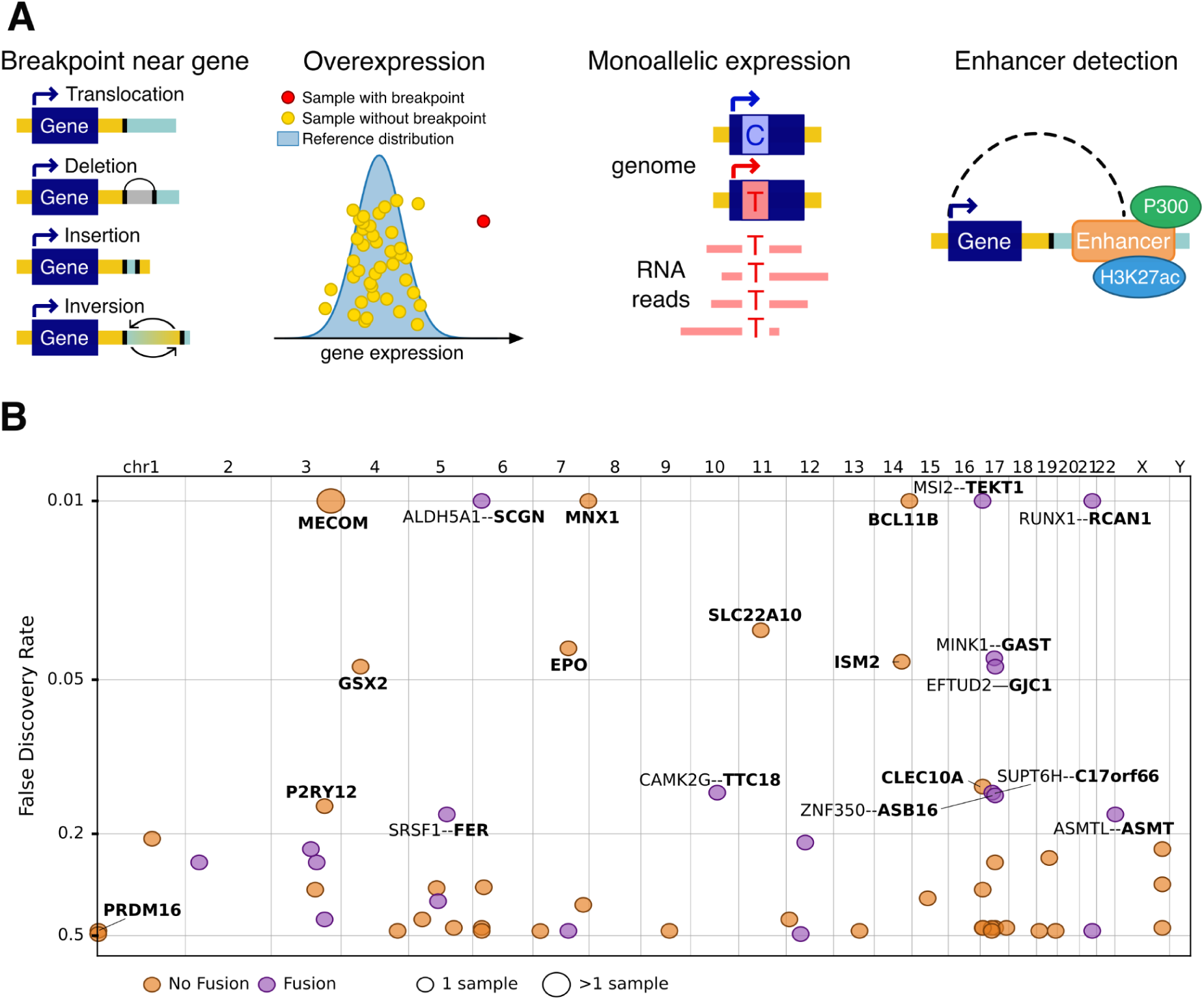
Detection of enhancer hijacking in 39 ckAML samples. **A.** Schematic representation of the main sources of information used by pyjacker: breakpoints, overexpression, monoallelic expression, and enhancers. **B.** Scatter plot of genes identified by pyjacker in 39 ckAML samples as being potentially activated by genomic rearrangements in one or more samples, where the x-axis shows the genomic location of the genes and the y-axis shows the FDR. Gene names for the enhancer hijacking candidates are written in bold, and if a fusion transcript was detected, the fusion partner is named.

### Putative enhancer hijacking events in 39 ckAML samples

We profiled 39 ckAML samples with WGS and RNA-seq. These samples were part of the ASTRAL-1 clinical trial which included older AML patients (29). These samples carried the alterations frequently found in ckAML (30), including bi-allelic *TP53* alterations (64%, N=25), del(7q) (69%, N=27), del(5q) (67%, N=26), and chromothripsis (43%, N=17) (Supplementary Fig. 1 and Supplementary Tables 4-8).

Pyjacker was applied to these 39 samples and detected 19 candidate genes with an FDR <20% (**Fig. 1B** and Supplementary Table 9). Among them were many of the genes which had previously been reported to be activated by enhancer hijacking in AML; including *MECOM* (in two samples), *MNX1* and *BCL11B*. In addition, pyjacker identified several genes that had not been reported before and which represent interesting candidate oncogenes to be verified in future studies. For 9 of the 19 genes, no fusion transcript was detected, suggesting enhancer hijacking as the underlying activation mechanism: *MECOM*, *MNX1*, *BCL11B*, *SLC22A10*, *EPO*, *ISM2*, *GSX2, CLEC10A* and *P2RY12*. In order to evaluate how recurrent the upregulation of these genes is in AML, we used data from the TCGA-LAML (1), BEAT-AML (31) and TARGET-AML (32) cohorts. We found that most of the genes identified by pyjacker were recurrently overexpressed in these other AML cohorts, albeit at low frequencies (Supplementary Fig. 2). However, some genes were not found overexpressed in these three other AML cohorts, which suggests either that their activation is a very rare event in AML, that they are false positives, or that their overexpression in our cohort was a passenger event of chromothriptic rearrangements. For example, the activations of *TEKT1* (in 16PB3075) and of *SLC22A10* (in 15KM20146) were due to complex rearrangements which also contained SVs within *TP53* (Supplementary Fig. 3). Thus, these rearrangements might have been selected for because of the *TP53* disruption rather than *TEKT1* or *SLC22A10* activation.

### Activation of *MECOM* and its homolog *PRDM16* by the *GATA2* enhancer

The only gene identified by pyjacker in more than one sample from this cohort was *MECOM*, found to be monoallelically overexpressed in two samples (Fig. 2A**-B** and Supplementary Fig. 4B). In both cases, the rearrangements were more complex than those found in samples with inv(3) or t(3;3) AML which are the most frequent rearrangements responsible for *MECOM* activation. One sample had chromothripsis on chromosome 3 (**Fig. 2C**), while the other one had several rearrangements between chromosome 3 and chromosome 14 (Supplementary Fig. 4A). Even though these rearrangements were very complex, they still resulted in the juxtaposition of *MECOM* to the *GATA2* enhancer (next to *RPN1*) (**Fig. 2D**), which is the same enhancer that activates *MECOM* in the more common inv(3) and t(3;3) (16). Interestingly, the *GATA2* enhancer was also reported by pyjacker to activate *PRDM16*, a homolog of *MECOM* (33), in another sample (16KM11270). This sample had a t(1;3)(p36;q21) translocation, which has been reported before as a rare event (33), and which juxtaposes *PRDM16* to the *GATA2* enhancer (**Fig. 2G**). Both *MECOM* (also known as *PRDM3*) and *PRDM16* are H3K9me1 methyltransferases (34), hence their overexpression could play a similar role in AML. Even though the expression of *PRDM16* was monoallelic (**Fig. 2F**), which is a strong indicator of activation by enhancer hijacking, the FDR reported by pyjacker was high (47%) because several samples without breakpoints near *PRDM16* had a higher expression than this sample (**Fig. 2E**). *MECOM* is also expressed in samples without breakpoints nearby (35), although to a lesser extent, suggesting an additional activation mechanism for *MECOM* and *PRDM16* besides enhancer hijacking.

**Figure 2.**
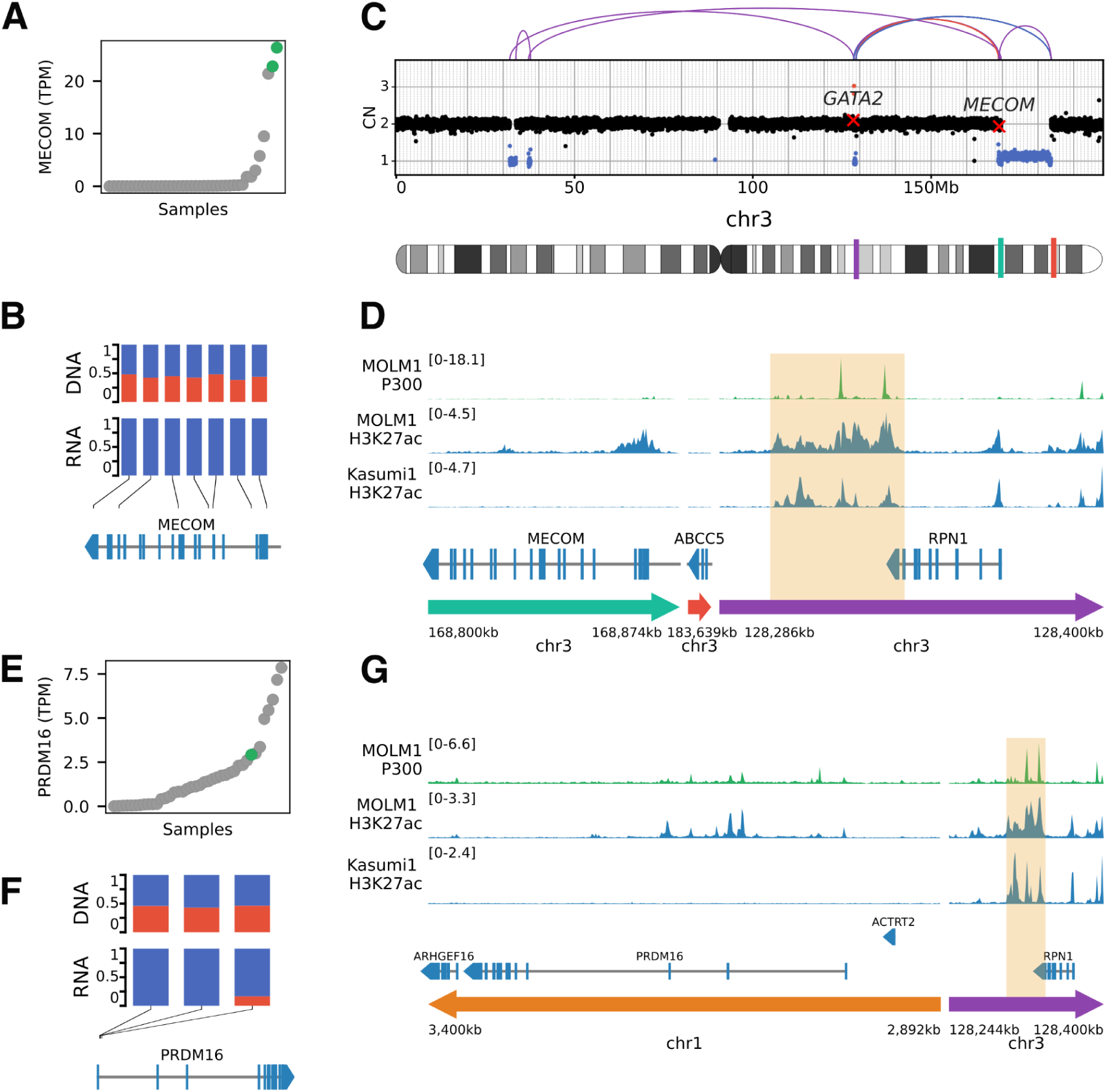
Activation of *MECOM* and its homolog *PRDM16* by the *GATA2* enhancer. **A**, **E**. Expression of *MECOM* and *PRDM16* in all samples, ranked by expression, where samples with a breakpoint near the corresponding gene are plotted in green. **B, F**. Variant allele frequencies in DNA and RNA for SNPs in *MECOM* and *PRDM16*, for samples 15PB19457 (with *MECOM* breakpoint) and 16KM11270 (with breakpoint near *PRDM16*). **C**. Copy numbers and SVs on chromosome 3 for sample 15PB19457. **D**. ChIP-seq tracks for P300 and H3K27ac in the myeloid cell lines MOLM-1 and Kasumi-1 in the region around *MECOM* for the rearranged chromosome of sample 15PB19457. **G**. ChIP-seq tracks for P300 and H3K27ac in the myeloid cell lines MOLM-1 and Kasumi-1 in the region around PRDM16 on the rearranged chromosome of sample 16KM11270.

### Aberrant *EPO* expression cooperates with *EPOR* amplification to drive acute erythroleukemia

Among the novel genes identified by pyjacker, an interesting candidate was *EPO*. To our knowledge, this gene has never been reported to be activated by enhancer hijacking in human leukemias, although it has been found to be overexpressed due to genomic rearrangements in a mouse model of erythroleukemia (36,37). *EPO* is not expressed in normal hematopoietic cells, but it is instead produced in the kidneys when blood oxygen levels are low, and it stimulates red blood cell proliferation by binding to its receptor (EPOR) and activating the JAK/STAT pathway (38–40). Since *EPO* promotes survival, proliferation and differentiation of erythroid progenitor cells (41), it may drive acute erythroleukemia (AEL), a rare subtype of AML enriched for complex karyotypes. In this ckAML cohort, the AEL sample 15KM18875 had high *EPO* expression (**Fig. 3A**). Although no samples from the TCGA-LAML, BEAT-AML and TARGET-AML cohorts expressed *EPO*, we found that among three AEL cohorts profiled with RNA-seq (42–44), one sample from each cohort expressed *EPO* (**Fig. 3B**), indicating that *EPO* expression is a rare but recurrent event in AEL. In sample 15KM18875, a 100 kb region on chromosome 7 around *EPO* was duplicated and fused with a region on chromosome 11 (**Fig. 3C**) such that an extrachromosomal circular DNA (eccDNA) was formed (**Fig. 3D**). eccDNAs are rather common in cancer, but they are often amplified, whereas sample 15KM18875 displayed an average copy number of less than one eccDNA per cell. This eccDNA is therefore subclonal, but it is unclear whether most cells have one copy, or whether a small percentage of cells contain numerous copies. The chromosome 11 portion of the eccDNA contains a putative enhancer with P300 and H3K27ac peaks in the leukemic cell line K562 with erythroid features (45), and this enhancer is likely responsible for the activation of *EPO* in this sample. In addition to high *EPO* expression, we also observed very high expression of the EPO receptor (*EPOR)* in 15KM18875 (**Fig. 3E**), which was due to a massive amplification of *EPOR* on chromosome 19 (**Fig. 3F**). Chromosome 19 harbored patterns of chromothripsis, as well as foldback inversions, suggesting that the amplifications were caused by breakage-fusion-bridge cycles (46). Amplification of *EPOR* has recently been reported as a recurrent driver event in AEL (44). High *EPOR* expression could make the cells very sensitive to *EPO*, thus increasing the fitness advantage provided by endogenous *EPO* expression by the leukemic cells. In both the Iacobucci et al. (42) and Fagnan et al. (43) cohorts, the sample with *EPO* expression also had outlier high *EPOR* expression, indicating that *EPO* is recurrently overexpressed together with *EPOR*.

**Figure 3.**
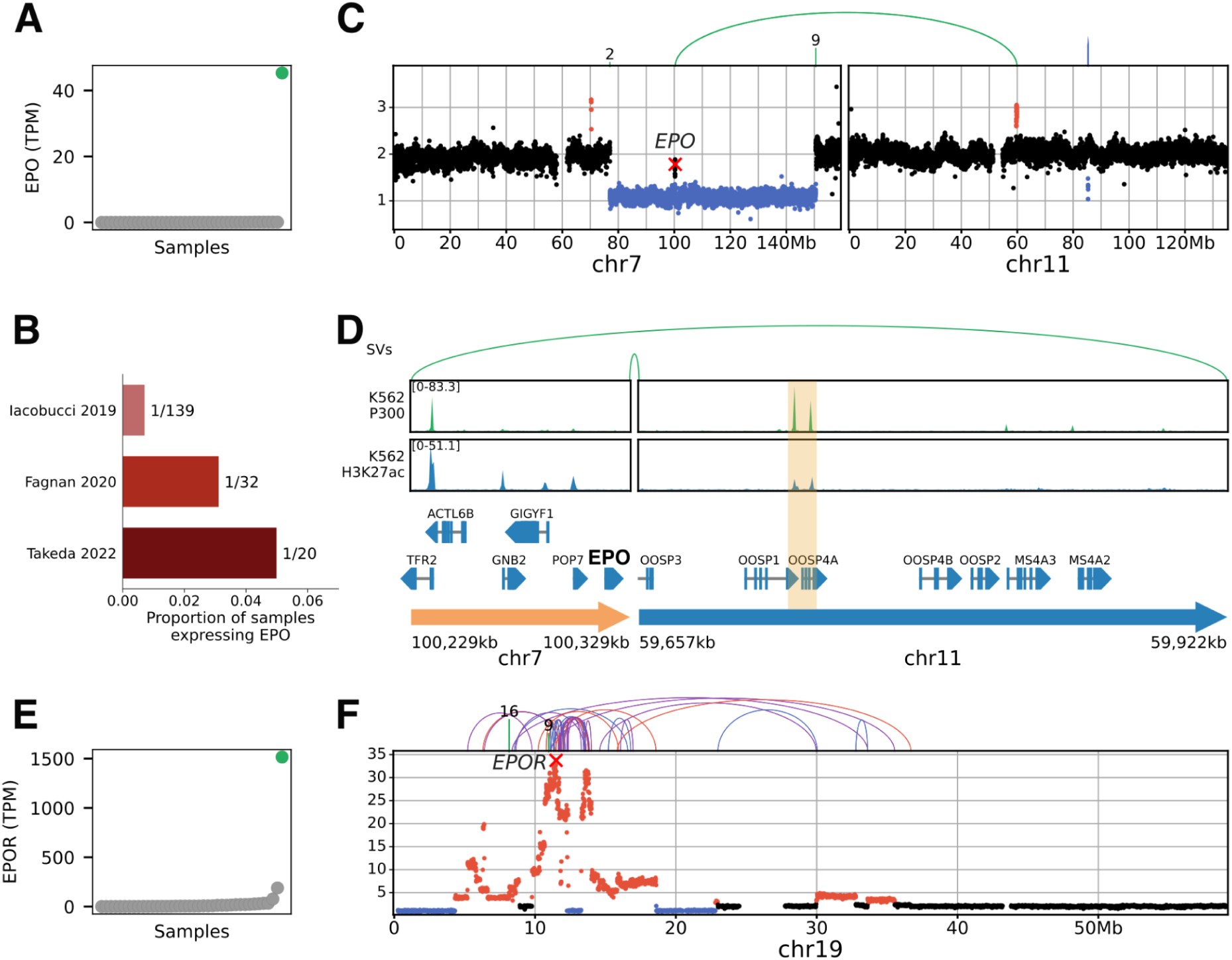
Aberrant *EPO* expression cooperates with *EPOR* amplification to drive acute erythroleukemia. **A**. *EPO* expression in all samples, with the sample 15KM18875 with *EPO* overexpression highlighted in green. **B.** Proportion of samples with *EPO* expression in three AEL cohorts profiled with RNA-seq (42–44). **C**. Copy numbers and SVs on chromosome 7 (containing *EPO*) and chromosome 11 in sample 15KM18875. **D.** 300 kb circular piece of DNA containing *EPO* and a putative enhancer (highlighted in yellow), with P300 and H3K27ac peaks in the erythroid cell line K562. **E**. *EPOR* expression in all samples, with sample 15KM18875 highlighted in green. **F**. Copy numbers and SVs on chromosome19 for sample 15KM18875.

### The homeobox genes *GSX2* and *MNX1* can be activated by atypical rearrangements

Among the top pyjacker hits were two homeobox genes, *GSX2* and *MNX1*, which were overexpressed in samples 16PB5693 and 15PB8708, respectively. Both samples have breakpoints near the respective genes, and in sample 15PB8708, heterozygous SNPs in *MNX1* confirmed monoallelic expression (**Fig. 4A-C**). Homeobox genes are often upregulated in AML (47), so the activation of homeobox genes by enhancer hijacking could be a driver event.

**Figure 4.**
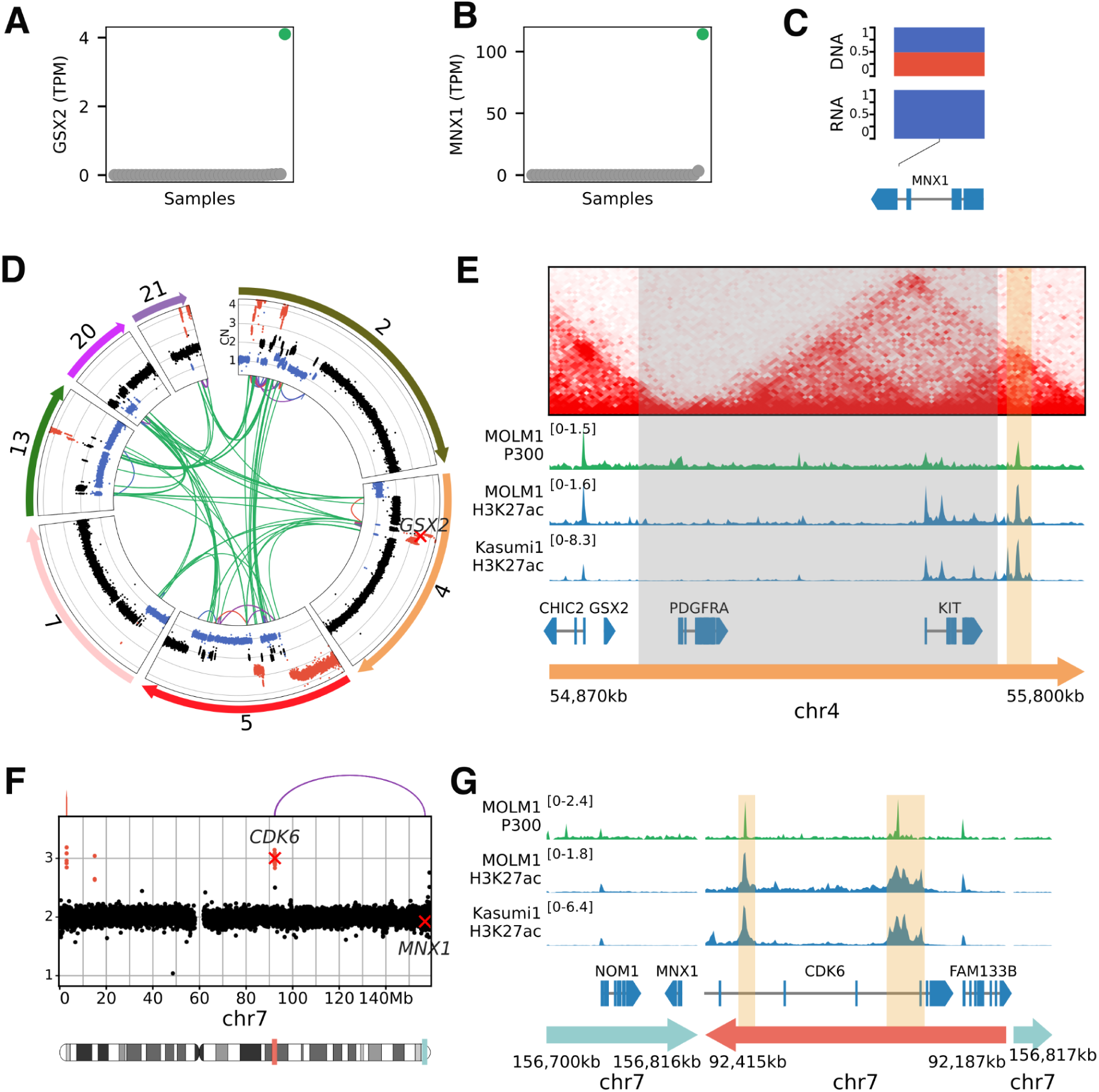
The homeobox genes *GSX2* and *MNX1* can be activated by atypical mechanisms. **A**. *GSX2* expression in all samples, with the sample 16PB5693 with *GSX2* expression highlighted in green. **B**. *MNX1* expression in all samples, with the sample 15PB8708 with *MNX1* overexpression highlighted in green. **C**. Variant allele frequencies in DNA and RNA for a SNP in *MNX1* in sample 15PB8708. **D**. Circos plot showing CNAs and SVs in sample 16PB5693, for the chromosomes involved in a chromothripsis event. **E**. HiC data from hematopoietic stem and progenitor cells (19) and ChIP-seq data from myeloid cell lines in the region around *GSX2*. The putative enhancer is highlighted in yellow and the region in gray is deleted in sample 16PB5693. **F**. Copy numbers and breakpoints on chromosome 7 for sample 15PB8708. **G**. ChIP-seq tracks for P300 and H3K27ac in the myeloid cell lines MOLM-1 and Kasumi-1 in the region around *MNX1*, on the rearranged chromosome of sample 15PB8708. Enhancers of the CDK6 region are highlighted in yellow.

Both *GSX2* and *MNX1* are known to be activated by rare but recurrent translocations to the *ETV6* locus; *GSX2* by t(4;12)(q11-q12;p13) in adult AML (48) and *MNX1* by t(7;12)(q36;p13) in pediatric AML (19). Here, however, *GSX2* and *MNX1* were activated by atypical mechanisms. Sample 16PB5693 was affected by a chromothripsis event involving multiple chromosomes, and several genomic segments, including *GSX2*, were amplified (**Fig. 4D**). In the wild-type state, the putative enhancer is located less than 1Mb away from *GSX2*, but in a different topologically-associating domain (TAD) (**Fig. 4E**). In sample 16PB5693, a deletion removed the TAD boundary, which likely enabled *GSX2* to interact with the enhancer. In addition to *GSX2* upregulation, the recurrent t(4;12) translocation frequently leads to *PDGFRA* activation and to an *ETV6::CHIC2* fusion transcript (49). Sample 16PB5693 only had *GSX2* expression without *PDGFRA* expression and without fusion transcript, suggesting that *GSX2* expression is the driving event. In sample 15PB8708, a 230 kb segment in the *CDK6* region, containing two putative enhancers, was duplicated and inserted next to *MNX1* (**Fig. 4F-G**). This hematopoietic super-enhancer has already been reported to be involved in enhancer hijacking events in AML, activating *BCL11B* (17) or *EVI1* (50). *MNX1* was expressed in a rather high proportion of the TCGA-LAML and BEAT-AML cohorts (2/179 and 17/707 samples with *MNX1* expression respectively), and the cytogenetic information often mentioned rearrangements of chromosome 7, including del(7)(q22q36), indicating that *MNX1* expression in these samples could be due to enhancer hijacking.

### *MNX1* is expressed in 1.4% of all AML cases, often with del(7)(q22q36)

To estimate the frequency of *MNX1* expression in AML cases, we performed an unbiased qRT-PCR screen of three different AML cohorts (Rotterdam, Ulm, Jena) (**Fig. 5A**). In a total of 2,293 cases across five cohorts [three qRT-PCR cohorts and public RNA-seq from TCGA-LAML (1) and BEAT-AML (31)], we estimated the frequency of *MNX1*-expressing samples to be 1.4% of all AML cases (Supplementary Table 10). We also screened del(7q) and ckAML cases and found a higher proportion of *MNX1*-expressing samples in these selected groups (8.70% in del(7q) and 2% in ckAML; Supplementary Table 10).

**Figure 5.**
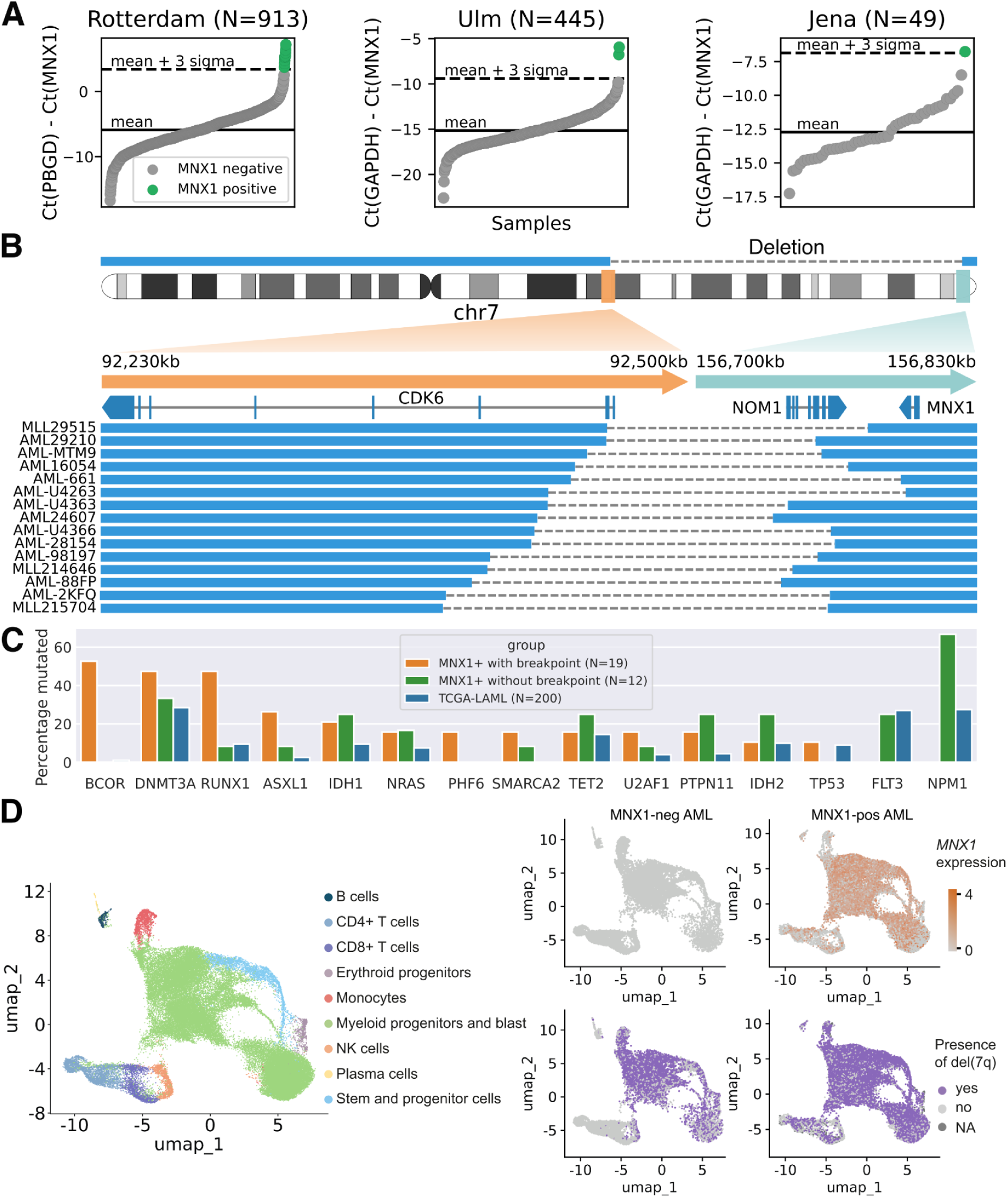
*MNX1* is expressed in 1.4% of all AML cases, often with del(7)(q22q36). **A**. qRT-PCR screen for *MNX1* expression in three AML cohorts (Rotterdam, Ulm, Jena). **B**. 15 *MNX1*-expressing samples with del(7)(q22q36) profiled with WGS, with a zoom-in around the breakpoints (hg19 reference). **C**. Percentage of samples with mutations in frequently mutated genes, for *MNX1*-positive samples with breakpoints near *MNX1*, *MNX1*-positive samples without breakpoints, and TCGA-LAML samples. **D**. ScRNA-seq analysis for *MNX1*-positive and control del(7q) AML samples. Left: UMAP showing cell type labels of 53,479 cells integrated across eight patients. Right: UMAP highlighting *MNX1* expression (top) and the presence of a del(7q) (bottom) as predicted for patients with del(7q) (n=4) and patients with del(7q) and *MNX1* activation (n=4).

We performed WGS on 23 *MNX1*-expressing samples, which we combined with WGS data of 8 samples provided by the Munich Leukemia Laboratory (MLL), resulting in a total of 31 *MNX1*-expressing samples profiled with WGS. Fifteen samples had a large del(7)(q22q36) starting within *CDK6* and ending before *MNX1* (**Fig. 5B**, Supplementary Table 11), indicating that *MNX1* could be activated by an enhancer in the *CDK6* region in those samples. Interestingly, this is the same region that is duplicated and inserted next to *MNX1* in sample 15PB8708 (**Fig. 4F-G**). Four samples had other rearrangements near *MNX1*, including a smaller del(7q) between the T-cell receptor beta locus and *MNX1* (Supplementary Fig. 5-6), which supports the notion that other enhancers apart from *CDK6* might activate *MNX1*. Indeed we had previously found a *MYB* enhancer in GDM-1 cells (27) and an *ETV6* enhancer in t(7;12)(q36;p13)^19^ pediatric AML to drive aberrant *MNX1* expression. Twelve samples had no rearrangements near *MNX1*, suggesting that *MNX1* may also be activated through other mechanisms.

Samples with *MNX1* rearrangements had a unique mutational spectrum with an absence of *NPM1* and *FLT3* mutations (0/19), as well as a very high frequency of *BCOR* mutations (10/19) which are usually rare in AML (2/200 in TCGA-LAML), although they have recently been reported to have a frequency of about 10% in AML with del(7q) (11) (**Fig. 5C** and Supplementary Table 12). *BCOR* mutations were accompanied by *BCORL1* (2/10) and *NCOR2* (1/10) mutations indicating a potential synergistic effect of multiple hits on this gene family. We also found *NCOR1* (1/9) and *NCOR2* (1/9) mutations in *BCOR-*wt cases, indicating that they might play a similar role as *BCOR* mutations. *MNX1*-expressing samples without breakpoints near *MNX1* did not share this mutational landscape. They had a particularly high frequency of mutations in *NPM1* (8/12), which could explain *MNX1* expression in these samples, since *NPM1* mutations can upregulate homeobox genes (51). *MNX1*, however, has not been shown to be in the *NPM1* gene signature in previous studies. In pediatric AML, *MNX1* can be expressed as a result of a translocation t(7;12), which very often co-occurs with trisomy 19 (19). However, trisomy 19 was not found in this cohort of adult *MNX1*-expressing samples.

We profiled 22/31 *MNX1*-positive samples with RNA-seq and found that they had a different gene expression signature, depending on whether the sample had a breakpoint near *MNX1* or not (Supplementary Fig. 7-8, Supplementary Table 13). *MNX1*-rearranged samples had a gene expression signature similar to t(7;12)(q36;p13) pediatric AML (19,52,53), with for example an upregulation of *AGR2*, *KRT72* and *KRT73.* Downregulated genes included several key cancer and hematopoiesis associated genes: *HLX*, *TFEC*, *GFI1*, *GAPT*, *SPRY2*, *TLE4*, *ACVR1B*, *BIK*, *EVI2B*, *PIK3CG*, *INPPL1 (SHIP2)*, *MYD88*, *MACC1*, *CSF3* and *CD177*. *MNX1*-non-rearranged samples had a different gene expression signature with a significant upregulation of *HOXA13*, *CCL1*, *CX3CR1* and a downregulation of *DLK1* and *DDIT4L*. *MNX1* expression was slightly lower than in *MNX1*-rearranged cases and some of the downregulated genes also showed intermediate levels in *MNX1*-non-rearranged samples.

Next we performed single-cell RNA sequencing (scRNA-seq) on eight AML samples (four *MNX1*-positive (*MNX1*+) and four *MNX1*-negative (*MNX1*-) with del(7q); Supplementary Fig. 9) to investigate the expression of *MNX1* and the presence of del(7q) at the single-cell level. We integrated scRNA-seq data for 53,479 cells across all patients and annotated the cell types by projecting the data onto a reference atlas (54) (**Fig. 5D**). We mainly captured myeloid progenitors and leukemic blasts, consistent with the disease phenotype. We observed that del(7q) was present in virtually all leukemic blasts across both groups (*MNX1*-and *MNX1*+), suggesting that this genomic alteration was an early event in leukemogenesis in these patients. In *MNX1*+ cases, *MNX1* was constitutively expressed in all blasts, indicating that cells with *MNX1* activation might have a proliferative advantage.

### Putative enhancers in the *CDK6* region interact with *MNX1* in del(7q) AML

Since most samples with *MNX1* activation have breakpoints in *CDK6*, we set out to identify the corresponding enhancer. To investigate whether *MNX1* may interact with the *CDK6* locus in selected del(7)(q22q36) samples, we performed circular chromosome conformation capture (4C) using a 5’ part of *MNX1* as viewpoint. In all three cases analyzed, 2KFQ, MTM9 and AML-661, we detected interactions between *MNX1* and the *CDK6* locus (**Fig. 6A**). We confirmed these interactions by reciprocal 4C using the *CDK6* locus as viewpoint (Supplementary Fig. 10). We further narrowed down the *CDK6*-derived enhancer to roughly 200 kb by combining the genomic information from the *CDK6* duplication of ckAML sample 15PB8708 and from the deletion margins of the del(7q) samples (**Fig. 6B**). Open chromatin profiling by ATAC-seq and enhancer mark profiling by ACT-seq in two patient samples and one PDX sample with del(7)(q22q36) revealed several enhancer candidates, two of which coincided with P300 and H3K27ac peaks in the MOLM-1 cell line (**Fig. 6B**). We considered the rightmost enhancer (chr7:92384001-92385000, hg19) located immediately at the deletion border as the strongest candidate and inserted it close to *MNX1* into one of the two chromosomes 7 of the induced pluripotent stem cell (iPSC) cell line ChiPSC22. Upon differentiation into hematopoietic stem and progenitor cells (HSPCs), the engineered, but not the wild-type cells showed *MNX1* expression, although at a significantly lower level than in patient samples (Supplementary Fig. 11). Therefore, this rightmost enhancer is not sufficient to induce the high *MNX1* expression observed in del(7)(q22q36) patients alone, and might require additional enhancers from this region. To recapitulate the genomic configuration of *MNX1* expressors with del(7q), we generated a heterozygous del(7)(q22q36) in the iPSC/HSPC model. However, del(7q) iPSCs could not be differentiated into HSPCs and therefore did not show *MNX1* activation, probably because haploinsufficiency of genes in the 60Mb deletion interfere with the differentiation (data not shown). Taken together, *MNX1* activation in del(7q)(q22q36) AML could be traced to a region of 200 kb including parts of *CDK6*. Identifying the precise location of the enhancer(s) will require future work.

**Figure 6.**
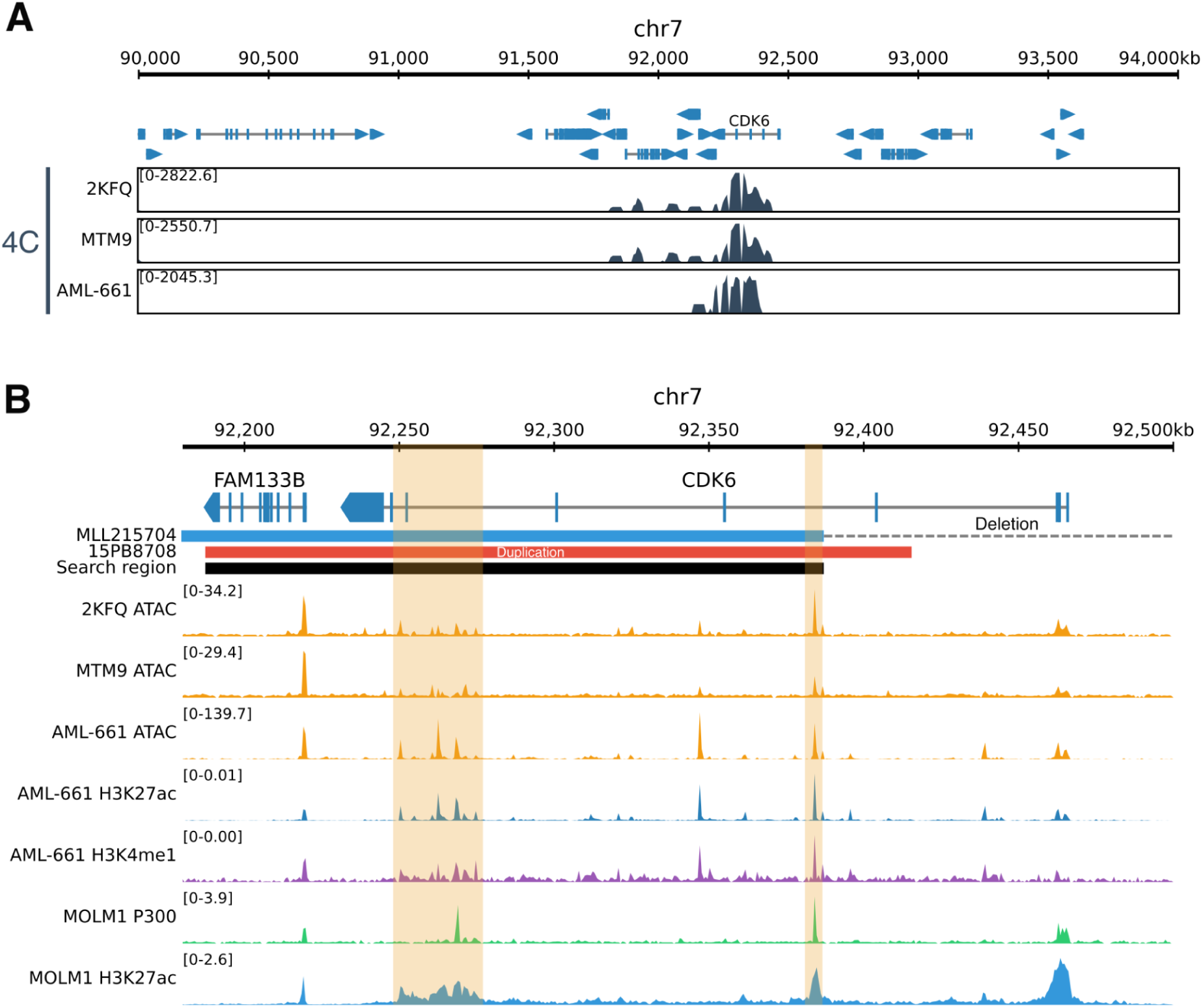
Putative enhancers in the *CDK6* region interact with *MNX1* in del(7q) AML. **A**. Chromatin interaction detected with 4C in the region around *CDK6* using *MNX1* as viewpoint, for three different del(7)(q22q36) samples. **B**. The 200 kb search region based on the enhancer duplication (sample 15PB8708) and the sample with the leftmost deletion (MLL215704), with tracks for enhancer marks: ATAC-seq in del(7q) samples MTM9 and 2KFQ, ATAC-seq and ACT-seq against H3K27ac and H3K4me1 in the PDX sample AML-661 derived from a del(7q) patient, and ChIP-seq against P300 and H3K27ac in the MOLM-1 cell line. The putative enhancers were highlighted.

### Knockdown of *MNX1* reduces tumor load of AML PDX cells *in vivo*

After having demonstrated that *MNX1* can be activated by enhancer hijacking in AML, we investigated whether *MNX1* plays a role in the maintenance of established leukemias. To approximate the clinical situation, we studied patients’ AML cells growing in mice, using PDX model AML-661 which harbors a del(7q) and expresses *MNX1*. Using lentiviruses, we stably expressed two different constructs in each cell, namely CRE-ERT2 in which CRE becomes activated by addition of Tamoxifen (TAM) and a CRE-inducible shRNA cassette in two different versions, for knockdown of either *MNX1* or a control gene. The two knockdown constructs were molecularly marked by different fluorochromes to distinguish the two populations by flow cytometry, before and after induction of the knockdown by TAM. In vivo experiments were performed in a competitive approach, injecting a mixture of cells with *MNX1* or control knockdown in a 1:1 ratio into the same mouse (**Fig. 7A**) (55). In the first, constitutive experiment, *MNX1* and control knockdowns were induced by TAM *in vitro* and before transplantation of PDX cells into mice (**Fig. 7A**). After a period of several weeks of leukemic growth in mice, cells with *MNX1* knockdown showed a pronounced disadvantage compared to cells with control knockdown in all organs studied (**Fig. 7B**), suggesting that lack of *MNX1* reduced fitness of PDX AML-661 cells *in vivo*. To distinguish the effect of *MNX1* knockdown on engraftment versus proliferation, a second experiment was performed where *MNX1* and control knockdowns were induced after the leukemic disease was readily established in mice, by systemic treatment of mice with TAM (**Fig. 7C**). Again, cells with *MNX1* knockdown had a remarkable disadvantage over control cells, most prominently in spleen and peripheral blood, indicating that *MNX1* knockdown reduced *in vivo* growth of AML-661 cells (**Fig. 7D**). As the effect was stronger in the first constitutive compared to the second inducible experiment, both biologic processes of cell engraftment and *in vivo* proliferation might rely on expression of *MNX1*.

**Figure 7.**
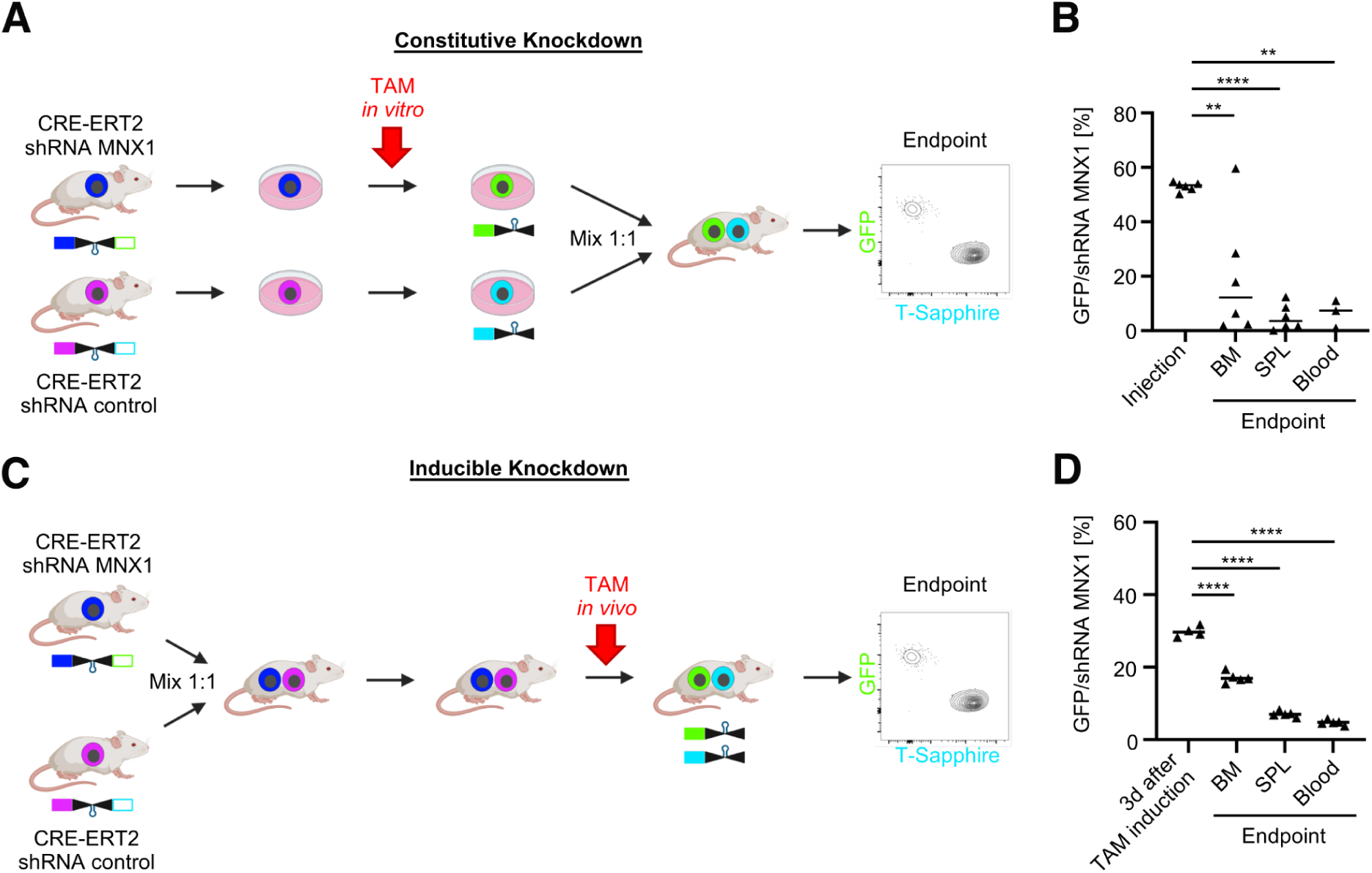
Knockdown of *MNX1* reduces tumor load of AML PDX cells in vivo. **A.** Scheme depicting the experimental setup of the in vivo constitutive experiment. AML-661 PDX cells expressing the cassettes for both CRE-ERT2 and the shRNA addressing *MNX1* or a control gene were amplified in mice. Fresh PDX cells were stimulated with Tamoxifen (TAM) to induce the knockdown *in vitro*. Cells with knockdown were enriched using flow cytometry gating on the respective fluorochrome markers GFP (knockdown of *MNX1*) and T-Sapphire (control knockdown). The two populations were mixed to a 1:1 ratio and injected into mice. The ratio between both populations was measured at advanced leukemic disease in different organs (endpoint). **B.** Results of the experiment described in **A** using 5 mice. **** *p* < 0.0001, ** *p* < 0.01 by paired t-test. **C.** Scheme depicting the experimental setup of the *in vivo* inducible experiment. The cell populations described in **A** were mixed in a 1:1 ratio and injected into 13 mice. 14 days after injection, 3 mice were sacrificed (N=3) to quality control the 1:1 ratio of the two cell populations using flow cytometry. Tamoxifen (TAM, 50 mg/kg) was orally administered to the 10 remaining mice. 5 mice were sacrificed 3 days later to measure the rate of shRNA induction by TAM. At an advanced stage of leukemia, the remaining 5 mice were sacrificed to determine the ratio between the control versus *MNX1* knockdown populations. **D.** Results of the experiment described in **C**. **** *p* < 0.0001 by unpaired t-test.

## Discussion

Sporadic reports have indicated enhancer hijacking as a mode of proto-oncogene activation in AML (16,17,19). Here, we developed pyjacker, a computational method for the systematic detection of enhancer hijacking events using WGS, RNA-seq data and enhancer information. Pyjacker is versatile and applicable to many cancer types, but here we focused on ckAML. In 39 ckAML samples, pyjacker detected 19 genes putatively activated by SVs in at least one sample with FDR<20%. This indicates the importance of enhancer hijacking in ckAML, although it is not as frequent as the most recurrent deletions in 5q and 7q. We found known genes activated by enhancer hijacking such as *MECOM, BCL11B* and *MNX1*, and identified multiple potential novel oncogenes in AML.

*GSX2* is a homeobox gene which is overexpressed in AML samples with the rare t(4;12)(q12;p13) translocation (48), but this translocation also often leads to overexpression of *PDGFRA* and fusions involving *ETV6*, the most frequent being *ETV6::CHIC2* (49). Here, we found a different rearrangement causing only *GSX2* overexpression without these additional effects, suggesting that activation of *GSX2* might be the driver event in the t(4;12) translocation and that understanding the role of *GSX2* in leukemogenesis could be important for therapeutic targeting.

*EPO* is another putative novel oncogene, activated by enhancer hijacking in a small fraction of AEL samples. *EPO* had already been found to be activated by structural rearrangements in a mouse model of erythroleukemia, resulting in growth factor independence (36,37). Here, we found one human AEL sample with *EPO* overexpression linked to a genomic rearrangement. Although *EPO* activation is rare, it appears to be recurrent in AEL, as we identified it in three additional cohorts (42–44), including a previously reported out-of-frame fusion transcript *YWHAE*::*EPO* which was probably selected for because it led to *EPO* upregulation (43). In addition, *EPO* overexpression seems to cooperate with amplifications of the gene coding for its receptor, a phenomenon recently described in AEL (44), since expression of *EPO* was found to co-occur with *EPOR* amplification.

Some newly identified genes were not found to be expressed in other cohorts, indicating that they may be very rare driver events, false positives, or passenger events which were selected for as part of a complex rearrangement. For example, both *TEKT1* and *SLC22A10* overexpression were caused by complex genomic rearrangements involving multiple chromosomes, which also disrupted *TP53*.

We focused validation experiments on *MNX1* since it was, among the top pyjacker hits, the second (behind *MECOM*) most recurrently expressed gene in other cohorts (1,31). We found that *MNX1* is expressed in 1.4% of all AML cases, often with del(7)(q22q36). Activation of *MNX1* with del(7q) had been reported before (56), and here we showed that the mechanism underlying the activation is a hijacking of a *CDK6* enhancer. Del(7q) is a recurrent event in AML and currently explained by haploinsufficiency of one or several genes, including *EZH2*, *KMT2C, KMT2E* and *CUX1* (11,13–15). Our findings show that, in addition to haploinsufficiency of the deleted genes, del(7q) can also lead to enhancer hijacking of *MNX1*. In one sample, a *CDK6* enhancer was duplicated and inserted next to *MNX1*, without deletion, which makes it very likely that *MNX1* activation is important for leukemogenesis, and not merely a passenger side effect of del(7q). *MNX1* upregulation had previously been observed in infant AML with t(7;12)(q36;p13) and was shown to transform fetal HSPCs in mice (19,57). Here, we showed that both constitutive and *in vivo* inducible knockdown of *MNX1* in competitive assays in an AML PDX model greatly reduced the fitness of the leukemic cells, which demonstrates that *MNX1* is a dependency gene in adult AML. However, only 8% of del(7q) AML cases have *MNX1* expression, so enhancer hijacking cannot explain all del(7q) cases and haploinsufficiency of genes in the deleted region remains the likely main consequence of del(7q). We found that this novel group of *MNX1*-rearranged adult AML samples have a unique mutational profile with a much higher rate of *BCOR* mutations (53%) than other AML samples (1%), and also higher than del(7q) AML (10%) (11). This differs from pediatric AML cases with t(7;12) which do not have these co-occurring *BCOR* mutations but instead frequently harbor trisomy 19 (19), an alteration that we did not detect in adult *MNX1*-rearranged cases. This new group of adult *MNX1*-rearranged patients had a gene expression signature that is similar to t(7;12) pediatric AML (52), suggesting that therapeutic strategies targeting *MNX1* could be jointly investigated for both pediatric and adult *MNX1*-rearranged AML cases. Suppression of key genes involved in hematological malignancies including *HLX*, *TFEC, GFI1*, *EVI2B*, *TLE4*, *MYD88*, all shared with pediatric AML, suggest a transcriptional repressor activity for *MNX1* in AML affecting cell proliferation and myeloid differentiation. As pediatric AML with *MNX1* activation has a different activation event, does not have chr7q deletions or *BCOR* mutations, and is seen in infants at a different developmental state, the overlap of dysregulated key genes strongly connects the observed gene dysregulation to *MNX1* activity and not to confounding factors. We also identified a novel group of *MNX1*-expressing cases without genomic rearrangements near *MNX1*, which do not share the gene expression signature of the *MNX1*-rearranged cases. The expression of *MNX1* in these samples remains unexplained, but we observed that they have a very high frequency of *NPM1* mutations (67%), which might be linked to *MNX1* expression, as *NPM1* mutations have been shown to upregulate homeobox genes (51). Notably, the differentially expressed genes in this group including *MNX1* and *HOXA13* do not overlap with previously described *NPM1*-associated gene signatures (58). Overall, this suggests the existence of undescribed modes of homeobox dysregulation in *NPM1* mutant AML, of which one could aberrantly activate *MNX1*.

Taken together, our data suggest that the numerous genomic rearrangements in ckAML often lead to enhancer hijacking, a molecular event that may have been previously underestimated compared with onco-fusions and CNAs. Understanding how the genes activated by this mechanism drive leukemia, or finding ways to stop this aberrant expression, could pave the way for personalized treatments targeting specific oncogenes.

## Methods

### Pyjacker details

#### Identification of “candidate samples” with breakpoints near a gene

Only genes whose expression is greater than 1 TPM (transcript per million) in at least one sample are considered. For each gene, pyjacker identifies “candidate samples” with a breakpoint near the gene, and which may therefore overexpress this gene because of the rearrangement. Since promoter-enhancer interactions occur within TADs, pyjacker selects samples which have a breakpoint in the same TAD as the gene. Any list of TADs can be provided, and in the present analysis we used TADs derived from publicly available HiC data from HSPCs (Supplementary Table 14) (19). To avoid missing events due to imprecise TAD boundaries, pyjacker extends the TADs by 80 kb on each side. If a list of TADs is not provided as input, pyjacker will instead consider all samples with breakpoints within a user-specified distance to the gene (1.5Mb by default). All “candidate samples” for a particular gene will be scored to test if these samples express this gene because of a structural rearrangement.

#### Overexpression score

If a gene is activated by enhancer hijacking in a sample, we expect this sample to have a higher expression for this gene, compared to “reference samples” which do not have breakpoints near the gene. In order to remove the effect of amplifications and to focus on genes activated by enhancer hijacking, the expression values in TPM are corrected for copy number, if CNA data is provided: the expression values are multiplied by 2/(copy number). The expression values are then log transformed: *log*(0. 5 + *E*). Then, pyjacker computes the mean μ and standard deviation σ of the gene expression in reference samples (which do not have breakpoints near the gene). For each candidate sample, pyjacker computes the number of standard deviations away its expression lies from the mean, where the standard deviation is increased in order to avoid extreme scores when all reference samples have the same expression: *t* = (*E* − µ)/(σ + 0. 3) where *E* is the expression of the gene in the candidate sample. This overexpression score is then transformed so that it is positive when the expression is more than two standard deviations above the mean and negative otherwise, and to avoid very high or very low overexpression scores which would have a disproportionate effect on the final score: if 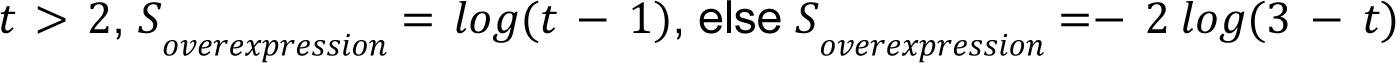.

#### Allele-specific expression (ASE) score

If a gene is activated by enhancer hijacking, we would expect only the allele on the rearranged chromosome to be expressed, resulting in monoallelic expression. For each gene and each sample, heterozygous SNPs are identified in the WGS data, and if there is coverage in the RNA-seq, the number of reference and alternative reads in the RNA-seq data are counted. For each SNP, pyjacker computes the log-likelihood ratio between monoallelic and bi-allelic expression. For monoallelic expression, we assume a mixture of two beta-binomial distributions for the allelic read counts, with means centered on 2% and 98% (to account for possible low expression from the other allele). For biallelic expression, we assume a beta binomial distribution centered on 50%. The log-likelihood ratios from all SNPs in the gene are then combined to get the allele-specific expression score, by averaging the log-likelihood ratios, but still giving a higher score if several SNPs are present. : 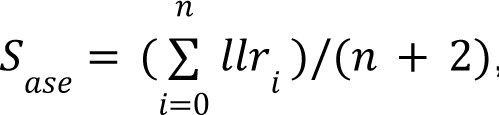, where *n* is the number of SNPs in the gene. This score is positive if the allelic information supports a monoallelic expression, negative if it supports a biallelic expression, and close to 0 if it is unclear. We note that if no heterozygous SNPs are present in a gene in a sample, the allele-specific expression score will be 0, but this does not preclude the gene from being identified by pyjacker, if the overexpression and enhancer scores are positive. The allele-specific expression score is set to 0 for genes with copy number lower than two or greater than four, for genes on sex chromosomes, and for imprinted genes (if a list of imprinted genes is provided as input). If allelic read counts are not provided as input, pyjacker can still be run and will in this case not use the allele-specific expression score, which will result in higher FDR.

#### Enhancer score

A genomic rearrangement is more likely to result in enhancer hijacking if it brings a strong enhancer close to the target gene. Pyjacker can optionally take as input a list of enhancers, scored for enrichment of enhancers marks by ROSE (59,60) (see section “Identification of myeloid enhancers” for the ChIP-seq data that we used in this study). The list of enhancers provided must be derived from the same cell type as the cancer samples studied. If no enhancer data is available, the enhancer score will be set to 0.

Pyjacker identifies all enhancers which, after the rearrangement, likely come to the same TAD as the gene. This is done by considering the position and orientation of the breakpoints, but each breakpoint is considered independently, which might miss some enhancers in case of complex rearrangements with clustered breakpoints. Enhancers are ranked according to their enrichment, and pyjacker computes the enhancer score by adding all scores, but putting more weight on the strongest enhancers: 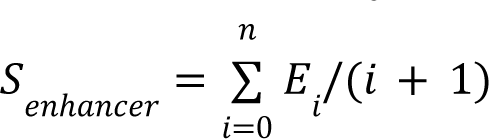 where *n* is the number of enhancers and *E_i_* is the enrichment for the *i*-th strongest enhancer.

#### Combined score

The overexpression, allele-specific expression and enhancer scores are then combined with a weighted sum. Pyjacker also penalizes if the gene is deleted in the sample, because rearrangements leading to enhancer hijacking should not delete the activated gene. This results in a score for each pair of (gene, candidate sample): 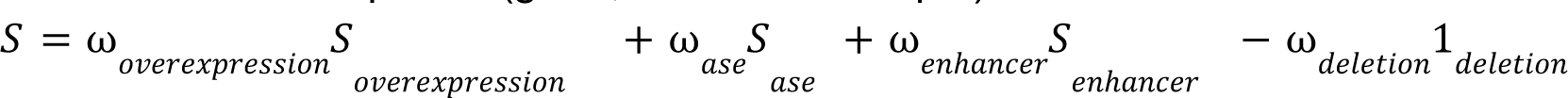 The weights can be set by the user, but their default values which should work well in all cases are 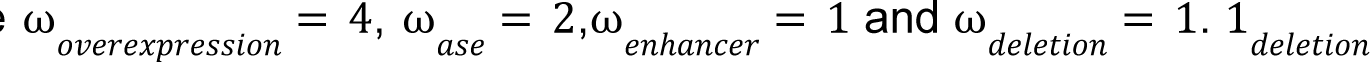 is 1 if the gene is deleted in the sample and 0 otherwise.

#### Aggregated gene score across samples

In order to give more weight to genes which are activated in multiple samples, pyjacker aggregates the scores from all samples for each gene: 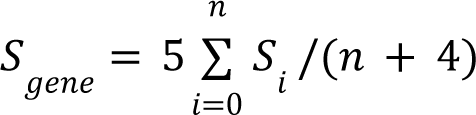 where *S_i_* is the score from sample *i*.

#### False discovery rate

The gene scores reflect how likely a gene is to be activated by structural rearrangements in the cohort studied, but the values are somewhat arbitrary. In order to get a more interpretable FDR, pyjacker computes a null distribution for these scores in the absence of enhancer hijacking. For each gene, the true “candidate samples” are excluded, and instead 1, 2 or 3 (number chosen randomly) random samples are chosen from the reference samples (without breakpoints near the gene) to be considered as candidate samples and scored. This results in a list of null scores, where only pairs of (gene, sample) without enhancer hijacking are used. The length of this list is equal to the number of genes (*n_genes_*), so to increase the size of the list (and thus get more precise p-values), this process is repeated *n_iter_* times (*n_iter_* = 50 by default), where each time different random samples are selected for each gene, resulting in a list of *n_iter_* * *n_genes_* null scores. This null distribution is used to compute an empirical p-value for each gene. Finally, the Benjamini-Hochberg correction is used to correct for multiple testing, which results in an FDR.

### AML cell lines used to test pyjacker

We tested pyjacker using 10 AML cell lines: THP-1, LAMA-84, MONOMAC-1, MV-4-11, HEL92.1.7, EOL-1, OCI-AML3, GDM-1, MOLM-1 and MUTZ-3. WGS and RNA-seq data for THP-1, LAMA-84, MONOMAC-1, MV-4-11, HEL92.1.7 and EOL-1 were retrieved for the Cancer Cell Line Encyclopedia (61). RNA-seq and WGS of GDM-1 were retrieved from GEO accession GSE221753 and SRA accession SRR23087016 (27). RNA-seq of OCI-AML3 was retrieved from GEO accession GSE209777 (62). WGS for OCI-AML3 and WGS and RNA-seq for MOLM-1 and MUTZ-3 were performed for this study and are available at https://www.ncbi.nlm.nih.gov/sra/PRJNA1140384.

### AML patient samples

The 39 ckAML samples were derived from a prospective clinical trial (NCT02348489) conducted in older, unfit patients with newly diagnosed AML (63). Data on targeted DNA sequencing of this cohort and in part of EPIC BeadChip arrays analysis were previously reported by Jahn et al. (29). For this study, we selected 39 ckAML samples (median age: 77 years), which had at least three CNAs detectable from the EPIC array data, and for which sufficient material was still available for further profiling.

### Generation and sequencing of whole genome sequencing data

DNA was isolated as previously described (19). The DNA was sequenced with NovaSeq 6000 S4, with read length of 2×150bp and a coverage of 50-70x for each sample. The WGS data was aligned to the GRCh37 reference genome using bwa-mem (arXiv:1303.3997v2 [q-bio.GN]). SVs were called with manta (64), CNAs were called with Control-FREEC (65) and SNVs with mutect2 (bioRxiv 10.1101/861054). Since no matched normal samples were available to identify somatic mutations, we only looked for SNVs in genes known to be recurrently mutated in AML, as previously described (19). Chromothripsis was determined using shatterseek (66), using as criterion at least 10 copy number switches in one chromosome. The WGS data processing, starting from the aligned bam files, was done using a nextflow workflow: https://github.com/CompEpigen/wf_WGS. All WGS plots were made using figeno (67).

### Generation and sequencing of RNA sequencing data

RNA was isolated as previously described (19). The RNA was sequenced with NovaSeq 6000 S2, with read length 2×101bp and 180-250 million reads per sample. The RNA-seq data was processed using the nf-core rnaseq workflow (68) v3.9, with alignment using STAR (69) and quantification using Salmon (70). Fusion transcripts were detected using Arriba (26). For allele-specific expression, we detected heterozygous SNPs in WGS data using HaplotypeCaller, and used GATK ASEReadCounter to get allele-specific read counts in RNA-seq data, at positions where a heterozygous SNP was found. Differential gene expression analysis was run using the deseq2 (71) package v1.42.0 with log fold change shrinkage applied by the ashr (72) algorithm v2.2-63. Batch correction was applied for the MLL cohort following the generation of vst-transformed gene expression values for single gene expression visualization. The TARGET pediatric AML RNA-seq dataset was downloaded from UCSC XENA and analyzed using the same approach as the adult AML cohort. For cases with multiple sample points, primary specimens were selected over recurrent samples. Bone marrow samples were preferentially used over blood-derived samples, yielding overall two unique cases with the t(7;12)(q36;p13) karyotype. The Balgobind et al. (52) pediatric AML cohort and its corresponding GEO GSE17855 Affymetrix U133 Plus 2.0 microarray dataset was analyzed using the Limma (73) package v3.58.1 using the empirical Bayes algorithm with default settings. Cases with unknown karyotype were not considered.

### Single-cell RNA sequencing of del(7q) AML patients

Single-cell RNA sequencing was performed for 8 AML samples: 4 *MNX1*-positive samples [3 with del(7q) and one with an alternative rearrangement] and 4 control *MNX1*-negative samples with del(7q). Cryopreserved samples from bone marrow and peripheral blood were thawed at 37°C for 2 min before transferring to a 50 mL tube. Cells were diluted by adding incremental 1:1 volumes of Gibco^TM^ DMEM/F12 media (Thermo Fisher Scientific) for five times with one-minute wait in between each step. Cells were centrifuged at 300 rcf for 5 min and resuspended in 2 mL PBS (Thermo Fischer Scientific) + 0.04% BSA (Milteny Biotec). Libraries were generated using 20,000 single cells as input to the Chromium Controller with the Chromium Next GEM Single Cell 3’ Kit v3.1 (10x Genomics). From the single-cell sequencing libraries, we generated between 632 and 803M (between 60,000 and 80,000 reads per cell) reads per sample using an Illumina NovaSeq 6000 S4 FlowCell. For processing (alignment to reference genome GRCh38, generation of count matrix) raw sequencing reads, cell ranger v7.1.0 was used. Subsequent analysis, including normalization (log-normalize), generation of a low dimensional representation, and cluster annotation was conducted using the Seurat v5 software package (74). Batch integration was performed with Canonical Correlation Analysis using Seurat’s IntegrateData function (75). For facilitating cluster annotation, we projected our data to the Triana et al. reference atlas (54) using scMap (76). We used numbat (77) for inferring copy number losses and gains from the single-cell transcriptomic data. A cell was annotated as having del(7q), if the probability of the deletion as returned by numbat was larger than 0.5.

### Identification of myeloid enhancers

We used public ChIP-seq data for H3K27ac and P300 from three myeloid cell lines: K562 (data from the ENCODE project (78), accessions ENCSR000AKP and ENCSR000EGE), MOLM-1 [data from array express accession E-MTAB-2224 (16)] and Kasumi-1 (data from GEO accession GSE167163; bioRxiv 10.1101/2022.09.14.507850). We used ROSE (59,60) to score and rank super enhancers, where transcription start sites were excluded. ROSE normally takes as input a single ChIP-seq experiment, but we found that the ranking was very variable depending on the dataset being used, so we used the six ChIP-seq datasets mentioned above and averaged the ROSE scores. The average ROSE scores were used as input to pyjacker, in order to compute the enhancer score.

### PDX model

Animal trials were performed in accordance with the current ethical standards of the official committee on animal experimentation (written approval by Regierung von Oberbayern; ROB-55.2Vet-2532.Vet_02-16-7 and ROB-55.2-2532.Vet_02-20-159). The PDX models AML-491 and AML-661 were established from an AML patient at first and second relapse. The cells harbored a del(7)(q21.13q36.3) and showed a positive *MNX1* expression. PDX cells were genetically modified as previously outlined in Zeller et al. (79). PDX cells were amplified in ten to 26-weeks-old male or female NOD.Cg-*Prkdc^scid^ Il2rg^tm1Wjl^*/SzJ (NSG) mice (The Jackson Laboratory, Bar Harbor, Maine, USA).

### Circular chromosome conformation capture (4C)

About two million cells per sample were used for circular chromosome conformation capture (4C) essentially according to van de Werken et al. (80). Two rounds of restriction digestion/T4 DNA ligation were applied, using *Bgl*II in combination with *Nla*III. In a first PCR step, second ligation products, inverse primers (Supplementary Table 15) and Q5 high fidelity enzyme (New England Biolabs, Frankfurt am Main, order no. M0491) were used with reaction conditions 98°C for 30 sec, 10 cycles with 98°C for 15 sec, 63°C, 57°C or 54°C, depending on the viewpoint, for 20 sec with 0.5°C touch-down per cycle, 72°C for 2 min, then 30 or 25 cycles with 98°C for 15 sec, 58°C, 52°C or 49°C, depending on the viewpoint, for 20 sec, 72°C for 2 min, finally followed by 72°C for 1 min. Purification of PCR products, generation of sequencing libraries and sequencing were done as described previously^25^.

### Antibody-guided Chromatin Tagmentation (ACT-seq)

Genome-wide targeting of histone modifications was done by ACT-seq according to Carter et al. (81) with some modifications using a self-prepared pA-Tn5ase protein (27), and using the antibodies listed in Supplementary Table 16.

### Assay for transposase-accessible chromatin by sequencing (ATAC-seq)

ATAC-seq was done essentially as described by Corces et al. (82) using about 50,000 cells and the Nextera DNA library prep kit (Illumina, Berlin, order no. 15028212). Libraries were generated and processed as described for ACT-seq, but cycling conditions were 98°C, 10 sec, 63°C, 30 sec, 72°C, 30 sec.

## 4C-seq, ACT-seq and ATAC-seq data analysis

4C-seq data processing and analysis was done with the pipe4C pipeline (83) using single reads starting with a *Bgl*II-site containing viewpoint primer; the pipe4C pipeline was applied with default parameters under R v3.6.2. ACT-seq and ATAC-seq data were analyzed as described previously^25^. Bigwig tracks were visualized using figeno (67).

### CRISPR/Cas9-mediated enhancer insertion

A 1 kb region (chr7:92384001-92385000, GRCh37/hg19) containing a putative enhancer was inserted upstream of the *MNX1* promoter (chr7:156816239, GRCh37/hg19) in ChiPSC22 (Takara Bio Europe) by CRISPR/Cas9 editing as previously described (84). In short, the cells were nucleofected with the Cas9 ribonucleoprotein complex and a homology directed repair (HDR) donor template containing the putative enhancer sequence and 200 bp homology arms on each site. The CRISPR RNA was designed using the Alt-R Custom Cas9 crRNA Design Tool (Integrated DNA Technologies) and the HDR donor template were ordered as dsDNA HDR Donor Blocks (Integrated DNA Technologies). Per 20 µL transfection, 500 ng of the HDR Donor Block were used. Clones with successful integration of the enhancer on one allele were selected by PCR, using the following primers: AAAAGGACATGGGGATGCGT and GAAGCTGATCTTCCCTGAGGTT. Two cell lines were validated using WGS. Cell lines were differentiated to hematopoietic stem and progenitor cells as previously described (84). RNA was isolated from HSPCs using the RNeasy Plus Mini Kit (Qiagen) and sequenced as described above.

### Competitive MNX1 knockdown *in vivo* assays

Tamoxifen-inducible shRNA constructs were generated as described in Carlet et al. (55) for two individual MNX1 shRNAs (76 & 82) and Renilla control shRNAs. Cre^ERT2^ and the shRNA cassettes were stably integrated into the AML-661 PDX model via lentiviral transduction. Cre^ERT2^/shMNX1-76, Cre^ERT2^/shMNX1-82, Cre^ERT2^/shRenilla-1 and Cre^ERT2^/shRenilla-2 transgenic cells were enriched with a BD FACSAria™ III Cell Sorter (BD Biosciences, Heidelberg) and serially transplanted into donor mice for amplification.

#### Constitutive Knockdown

Transgenic AML PDX cells were isolated from bone marrow of donor mice and cultured in StemPro-34 medium (Thermo Fisher Scientific) with Pen/Strep, L-Glutamine (both Gibco), 10 ng/ml hrFLT3L (R&D Systems), 10 ng/ml hrSCF, 10 ng/ml hrTPO, and 10 ng/ml hrIL3 (all Peprotech) (85) at a density of 10^6^ cells/ml at 37 °C, 5% CO_2_. For *ex vivo* flipping of the shRNA cassettes, the cells were treated using 200nM (Z)-4-Hydroxytamoxifen (Sigma Aldrich, St. Louis, USA, order no. H7904). This induces flipping of the shRNA cassette, which leads to the expression of the respective shRNA and a switch of the expressed fluorochrome from mTagBFP to eGFP and from iRFP720 to T-Sapphire, respectively. Cells harboring the flipped cassette were enriched via FACS. MNX1 shRNA and Renilla control shRNA expressing cells were mixed in a 1:1 ratio and injected into three mice per MNX1 shRNA via tail vein injection (1*10^6^ cells per population 2*10^6^ per mouse). The individual input mixes were measured using flow cytometry for each animal before injection as an input sample. Outgrowth of tumor cells was monitored by repeated blood samplings and staining for hCD33+ cells. At an advanced stage of leukemia (hCD33+ cells > 60%), mice were sacrificed and PDX cells were isolated from the bone marrow, spleen and blood. The ratio of the two cell populations was measured with flow cytometry as output samples, and the data was analyzed using Prism 10 (GraphPad Prism, La Jolla, USA).

#### Inducible Knockdown

*In vivo* induction of the MNX1 shRNA expression was performed according to Carlet et al. (55). Transgenic AML PDX cells were isolated from bone marrow of donor mice. Cre^ERT2^/shMNX1 and Cre^ERT2^/shRenilla transgenic cells were mixed in a 1:1 ratio and injected into mice via tail vein injection (N = 13; 1×10^6^ cells per population and mouse). 50 mg/kg tamoxifen (Sigma Aldrich, St. Louis, USA, order no. T5648) was administered once 14 days post-transplantation via oral gavage as previously described. Mice were sacrificed on the day of TAM administration without receiving TAM, three days after TAM administration, and at an advanced stage of leukemia (hCD33+ cells > 60%). The ratio of the two flipped cell populations was measured using flow cytometry and the data were analyzed using Prism 10 (GraphPad Prism, La Jolla, USA).

## Data availability

WGS and RNA-seq data of patient samples will be uploaded to EGA. WGS of the cell line OCI-AML3 and WGS and RNA-seq of the cell lines MOLM-1 and MUTZ-3 were uploaded to the SRA under project <u>PRJNA1140384</u>.

## Code availability

The source code for pyjacker is available at https://github.com/CompEpigen/pyjacker. The nextflow workflow used to prepare pyjacker’s inputs, starting from bam files, is available at https://github.com/CompEpigen/wf_WGS.

## Acknowledgements

We thank the Genomics and Proteomics Core Facility, the Omics IT and Data Management Core Facility and the Single Cell Open Lab of the DKFZ Heidelberg. We thank Ilaria Iacobucci and Charles Mullighan (St. Jude Children’s Research Hospital, Memphis) for sharing their AEL RNA-seq dataset. We thank June Takeda and Seishi Ogawa (Kyoto University) for sharing their AEL RNA-seq dataset.

## Funding

This work was supported in part by the German Funding Agency (DFG) through SPP1463 (to DBL & CP) and SFB 1074 (to HD, KD, LB, CP), the Carreras Foundation (CP), the Helmholtz Foundation, the Heidelberg Center for Personalized Oncology (HIPO) and the NCT Personalized Oncology Program (NCT-POP; project #HIPO-030 to DBL and CP). F. Heidel was supported by the Thuringian state program ProExzellenz (RegenerAging - FSU-I-03/14) and through project grants of the German Research Council (DFG) HE6233/4-2, project number 320028127, HE6233/9-1 project number 453491106 and HE6233/10-1 project number 505859092. M. Scherer is supported through a postdoctoral fellowship by the Dr. Rurainski Foundation. M. Schönung is supported by the Joachim Herz Foundation (Add-on Fellowships for Interdisciplinary Life Science).

## Authors’ disclosures

UHT is currently employed at Oxford Nanopore Technologies. EJ is currently employed at AstraZeneca. LB has received honoraria from AbbVie, Amgen, Astellas, BristolMyers Squibb, Celgene, Daiichi Sankyo, Gilead, Hexal, Janssen, Jazz Pharmaceuticals, Menarini, Novartis, Pfizer, Roche, and Sanofi, as well as research support from Bayer and Jazz Pharmaceuticals. DBL received honoraria from Infectopharm GmbH. All other authors declared no conflict of interest.

## References

1. TCGA. Genomic and Epigenomic Landscapes of Adult De Novo Acute Myeloid Leukemia. N Engl J Med. 2013;368:2059–74

2. Papaemmanuil E, Gerstung M, Bullinger L, Gaidzik VI, Paschka P, Roberts ND, et al. Genomic Classification and Prognosis in Acute Myeloid Leukemia. N Engl J Med. 2016;374:2209–21

3. Arber DA, Orazi A, Hasserjian RP, Borowitz MJ, Calvo KR, Kvasnicka H-M, et al. International Consensus Classification of Myeloid Neoplasms and Acute Leukemias: integrating morphologic, clinical, and genomic data. Blood. 2022;140:1200–28

4. Mrózek K. Cytogenetic, Molecular Genetic, and Clinical Characteristics of Acute Myeloid Leukemia With a Complex Karyotype. Semin Oncol. 2008;35:365–77

5. Korbel JO, Campbell PJ. Criteria for Inference of Chromothripsis in Cancer Genomes. Cell. 2013;152:1226–36

6. Rode A, Maass KK, Willmund KV, Lichter P, Ernst A. Chromothripsis in cancer cells: An update. Int J Cancer. 2016;138:2322–33

7. Fontana MC, Marconi G, Feenstra JDM, Fonzi E, Papayannidis C, Ghelli Luserna di Rorá A, et al. Chromothripsis in acute myeloid leukemia: biological features and impact on survival. Leukemia. 2018;32:1609–20

8. Schoch C, Haferlach T, Bursch S, Gerstner D, Schnittger S, Dugas M, et al. Loss of genetic material is more common than gain in acute myeloid leukemia with complex aberrant karyotype: A detailed analysis of 125 cases using conventional chromosome analysis and fluorescence in situ hybridization including 24-color FISH. Genes Chromosomes Cancer. 2002;35:20–9

9. Rücker FG, Bullinger L, Schwaenen C, Lipka DB, Wessendorf S, Fröhling S, et al. Disclosure of Candidate Genes in Acute Myeloid Leukemia With Complex Karyotypes Using Microarray-Based Molecular Characterization. J Clin Oncol. 2006;24:3887–94

10. Schoch C, Kern W, Kohlmann A, Hiddemann W, Schnittger S, Haferlach T. Acute myeloid leukemia with a complex aberrant karyotype is a distinct biological entity characterized by genomic imbalances and a specific gene expression profile. Genes Chromosomes Cancer. 2005;43:227–38

11. Halik A, Tilgner M, Silva P, Estrada N, Altwasser R, Jahn E, et al. Genomic characterization of AML with aberrations of chromosome 7: a multinational cohort of 519 patients. J Hematol OncolJ Hematol Oncol. 2024;17:70

12. Greenberg P, Cox C, LeBeau MM, Fenaux P, Morel P, Sanz G, et al. International Scoring System for Evaluating Prognosis in Myelodysplastic Syndromes. Blood. 1997;89:2079–88

13. Inaba T, Honda H, Matsui H. The enigma of monosomy 7. Blood. 2018;131:2891–8

14. McNerney ME, Brown CD, Wang X, Bartom ET, Karmakar S, Bandlamudi C, et al. CUX1 is a haploinsufficient tumor suppressor gene on chromosome 7 frequently inactivated in acute myeloid leukemia. Blood. 2013;121:975–83

15. Chen C, Liu Y, Rappaport AR, Kitzing T, Schultz N, Zhao Z, et al. MLL3 Is a Haploinsufficient 7q Tumor Suppressor in Acute Myeloid Leukemia. Cancer Cell. 2014;25:652–65

16. Gröschel S, Sanders MA, Hoogenboezem R, de Wit E, Bouwman BAM, Erpelinck C, et al. A Single Oncogenic Enhancer Rearrangement Causes Concomitant EVI1 and GATA2 Deregulation in Leukemia. Cell. 2014;157:369–81

17. Montefiori LE, Bendig S, Gu Z, Chen X, Pölönen P, Ma X, et al. Enhancer Hijacking Drives Oncogenic BCL11B Expression in Lineage-Ambiguous Stem Cell Leukemia. Cancer Discov. 2021;11:2846–67

18. von Bergh ARM, van Drunen E, van Wering ER, van Zutven LJCM, Hainmann I, Lönnerholm G, et al. High incidence of t(7;12)(q36;p13) in infant AML but not in infant ALL, with a dismal outcome and ectopic expression of HLXB9. Genes Chromosomes Cancer. 2006;45:731–9

19. Weichenhan D, Riedel A, Sollier E, Toprak UH, Hey J, Breuer KH, et al. Altered enhancer-promoter interaction leads to MNX1 expression in pediatric acute myeloid leukemia with t(7;12)(q36;p13). Blood Adv. 2024;bloodadvances.2023012161

20. Weischenfeldt J, Dubash T, Drainas AP, Mardin BR, Chen Y, Stütz AM, et al. Pan-cancer analysis of somatic copy-number alterations implicates IRS4 and IGF2 in enhancer hijacking. Nat Genet. 2017;49:65–74

21. Zhang Y, Chen F, Creighton CJ. SVExpress: identifying gene features altered recurrently in expression with nearby structural variant breakpoints. BMC Bioinformatics. 2021;22:135

22. Yu A, Yesilkanal AE, Thakur A, Wang F, Yang Y, Phillips W, et al. HYENA detects oncogenes activated by distal enhancers in cancer. Nucleic Acids Res. 2024;52:e77

23. Liu Y, Li C, Shen S, Chen X, Szlachta K, Edmonson MN, et al. Discovery of regulatory noncoding variants in individual cancer genomes by using cis-X. Nat Genet. 2020;52:811–8

24. Wang X, Xu J, Zhang B, Hou Y, Song F, Lyu H, et al. Genome-wide detection of enhancer-hijacking events from chromatin interaction data in rearranged genomes. Nat Methods. 2021;18:661–8

25. Haas BJ, Dobin A, Li B, Stransky N, Pochet N, Regev A. Accuracy assessment of fusion transcript detection via read-mapping and de novo fusion transcript assembly-based methods. Genome Biol. 2019;20:213

26. Uhrig S, Ellermann J, Walther T, Burkhardt P, Fröhlich M, Hutter B, et al. Accurate and efficient detection of gene fusions from RNA sequencing data. Genome Res. 2021;31:448–60

27. Weichenhan D, Riedel A, Meinen C, Basic A, Toth R, Bähr M, et al. Translocation t(6;7) in AML-M4 cell line GDM-1 results in MNX1 activation through enhancer-hijacking. Leukemia. 2023;1–4

28. Riedel SS, Lu C, Xie HM, Nestler K, Vermunt MW, Lenard A, et al. Intrinsically disordered Meningioma-1 stabilizes the BAF complex to cause AML. Mol Cell. 2021;81:2332–2348.e9

29. Jahn E, Saadati M, Fenaux P, Gobbi M, Roboz GJ, Bullinger L, et al. Clinical impact of the genomic landscape and leukemogenic trajectories in non-intensively treated elderly acute myeloid leukemia patients. Leukemia. 2023;37:2187–96

30. Rücker FG, Schlenk RF, Bullinger L, Kayser S, Teleanu V, Kett H, et al. TP53 alterations in acute myeloid leukemia with complex karyotype correlate with specific copy number alterations, monosomal karyotype, and dismal outcome. Blood. 2012;119:2114–21

31. Bottomly D, Long N, Schultz AR, Kurtz SE, Tognon CE, Johnson K, et al. Integrative analysis of drug response and clinical outcome in acute myeloid leukemia. Cancer Cell. 2022;40:850–864.e9

32. Bolouri H, Farrar JE, Triche T, Ries RE, Lim EL, Alonzo TA, et al. The molecular landscape of pediatric acute myeloid leukemia reveals recurrent structural alterations and age-specific mutational interactions. Nat Med. 2018;24:103–12

33. Mochizuki N, Shimizu S, Nagasawa T, Tanaka H, Taniwaki M, Yokota J, et al. A novel gene, *MEL1*, mapped to 1p36.3 is highly homologous to the *MDS1/EVI1* gene and is transcriptionally activated in t(1;3)(p36;q21)-positive leukemia cells. Blood. 2000;96:3209–14

34. Pinheiro I, Margueron R, Shukeir N, Eisold M, Fritzsch C, Richter FM, et al. Prdm3 and Prdm16 are H3K9me1 Methyltransferases Required for Mammalian Heterochromatin Integrity. Cell. 2012;150:948–60

35. Gröschel S, Lugthart S, Schlenk RF, Valk PJM, Eiwen K, Goudswaard C, et al. High EVI1 Expression Predicts Outcome in Younger Adult Patients With Acute Myeloid Leukemia and Is Associated With Distinct Cytogenetic Abnormalities. J Clin Oncol. 2010;28:2101–7

36. Howard JC, Berger L, Bani MR, Hawley RG, Ben-David Y. Activation of the erythropoietin gene in the majority of F-MuLV-induced erythroleukemias results in growth factor independence and enhanced tumorigenicity. Oncogene. 1996;12:1405–15

37. Chrétien S, Duprez V, Maouche L, Gisselbrecht S, Mayeux P, Lacombe C. Abnormal erythropoietin (Epo) gene expression in the murine erythroleukemia IW32 cells results from a rearrangement between the G-protein β2 subunit gene and the Epo gene. Oncogene. 1997;15:1995–9

38. Klingmüller U, Bergelson S, Hsiao JG, Lodish HF. Multiple tyrosine residues in the cytosolic domain of the erythropoietin receptor promote activation of STAT5. Proc Natl Acad Sci. 1996;93:8324–8

39. Richmond TD, Chohan M, Barber DL. Turning cells red: signal transduction mediated by erythropoietin. Trends Cell Biol. 2005;15:146–55

40. Jayavelu AK, Schnöder TM, Perner F, Herzog C, Meiler A, Krishnamoorthy G, et al. Splicing factor YBX1 mediates persistence of JAK2-mutated neoplasms. Nature. 2020;588:157–63

41. Perner F, Perner C, Ernst T, Heidel FH. Roles of JAK2 in Aging, Inflammation, Hematopoiesis and Malignant Transformation. Cells. 2019;8:854

42. Iacobucci I, Wen J, Meggendorfer M, Choi JK, Shi L, Pounds SB, et al. Genomic subtyping and therapeutic targeting of acute erythroleukemia. Nat Genet. 2019;51:694–704

43. Fagnan A, Bagger FO, Piqué-Borràs M-R, Ignacimouttou C, Caulier A, Lopez CK, et al. Human erythroleukemia genetics and transcriptomes identify master transcription factors as functional disease drivers. Blood. 2020;136:698–714

44. Takeda J, Yoshida K, Nakagawa MM, Nannya Y, Yoda A, Saiki R, et al. Amplified EPOR/JAK2 Genes Define a Unique Subtype of Acute Erythroid Leukemia. Blood Cancer Discov. 2022;3:410–27

45. Andersson LC, Nilsson K, Gahmberg CG. K562—A human erythroleukemic cell line. Int J Cancer. 1979;23:143–7

46. Lo AWI, Sabatier L, Fouladi B, Pottier G, Ricoul M, Murnane JP. DNA Amplification by Breakage/Fusion/Bridge Cycles Initiated by Spontaneous Telomere Loss in a Human Cancer Cell Line. Neoplasia N Y N. 2002;4:531–8

47. Khan I, Amin MA, Eklund EA, Gartel AL. Regulation of HOX gene expression in AML. Blood Cancer J. 2024;14:1–6

48. Cools J, Mentens N, Odero MD, Peeters P, Wlodarska I, Delforge M, et al. Evidence for position effects as a variantETV6-mediated leukemogenic mechanism in myeloid leukemias with a t(4;12)(q11-q12;p13) or t(5;12)(q31;p13). Blood. 2002;99:1776–84

49. Müller-Jochim A, Meggendorfer M, Walter W, Haferlach T, Kern W, Haferlach C. AML with t(4;12)(q12;p13): A Detailed Genomic and Transcriptomic Analysis Reveals Genomic Breakpoint Heterogeneity, Absence of Pdgfra Fusion Transcripts and Presence of Pdgfra Overexpression in a Subset of Cases. Blood. 2023;142:6014

50. Ottema S, Mulet-Lazaro R, Beverloo HB, Erpelinck C, van Herk S, van der Helm R, et al. Atypical 3q26/MECOM rearrangements genocopy inv(3)/t(3;3) in acute myeloid leukemia. Blood. 2020;136:224–34

51. Brunetti L, Gundry MC, Sorcini D, Guzman AG, Huang Y-H, Ramabadran R, et al. Mutant NPM1 maintains the leukemic state through HOX expression. Cancer Cell. 2018;34:499–512.e9

52. Balgobind BV, Heuvel-Eibrink MMV den, Menezes RXD, Reinhardt D, Hollink IHIM, Arentsen-Peters STJCM, et al. Evaluation of gene expression signatures predictive of cytogenetic and molecular subtypes of pediatric acute myeloid leukemia. Haematologica. 2011;96:221–30

53. Ragusa D, Cicirò Y, Federico C, Saccone S, Bruno F, Saeedi R, et al. Engineered model of t(7;12)(q36;p13) AML recapitulates patient-specific features and gene expression profiles. Oncogenesis. 2022;11:1–7

54. Triana S, Vonficht D, Jopp-Saile L, Raffel S, Lutz R, Leonce D, et al. Single-cell proteo-genomic reference maps of the hematopoietic system enable the purification and massive profiling of precisely defined cell states. Nat Immunol. 2021;22:1577–89

55. Carlet M, Völse K, Vergalli J, Becker M, Herold T, Arner A, et al. In vivo inducible reverse genetics in patients’ tumors to identify individual therapeutic targets. Nat Commun. 2021;12:5655

56. Federico C, Owoka T, Ragusa D, Sturiale V, Caponnetto D, Leotta CG, et al. Deletions of chromosome 7q affect nuclear organization and HLXB9 gene expression in hematological disorders. Cancers. 2019;11:585

57. Waraky A, Östlund A, Nilsson T, Weichenhan D, Lutsik P, Bähr M, et al. Aberrant MNX1 expression associated with t(7;12)(q36;p13) pediatric acute myeloid leukemia induces the disease through altering histone methylation. Haematologica. 2024;109:725–39

58. Verhaak RGW, Goudswaard CS, van Putten W, Bijl MA, Sanders MA, Hugens W, et al. Mutations in nucleophosmin (NPM1) in acute myeloid leukemia (AML): association with other gene abnormalities and previously established gene expression signatures and their favorable prognostic significance. Blood. 2005;106:3747–54

59. Lovén J, Hoke HA, Lin CY, Lau A, Orlando DA, Vakoc CR, et al. Selective Inhibition of Tumor Oncogenes by Disruption of Super-Enhancers. Cell. 2013;153:320–34

60. Whyte WA, Orlando DA, Hnisz D, Abraham BJ, Lin CY, Kagey MH, et al. Master Transcription Factors and Mediator Establish Super-Enhancers at Key Cell Identity Genes. Cell. 2013;153:307–19

61. Ghandi M, Huang FW, Jané-Valbuena J, Kryukov GV, Lo CC, McDonald ER, et al. Next-generation characterization of the Cancer Cell Line Encyclopedia. Nature. 2019;569:503–8

62. Goyal A, Bauer J, Hey J, Papageorgiou DN, Stepanova E, Daskalakis M, et al. DNMT and HDAC inhibition induces immunogenic neoantigens from human endogenous retroviral element-derived transcripts. Nat Commun. 2023;14:6731

63. Fenaux P, Gobbi M, Kropf PL, Issa J-PJ, Roboz GJ, Mayer J, et al. Guadecitabine vs treatment choice in newly diagnosed acute myeloid leukemia: a global phase 3 randomized study. Blood Adv. 2023;7:5027–37

64. Chen X, Schulz-Trieglaff O, Shaw R, Barnes B, Schlesinger F, Källberg M, et al. Manta: rapid detection of structural variants and indels for germline and cancer sequencing applications. Bioinformatics. 2016;32:1220–2

65. Boeva V, Popova T, Bleakley K, Chiche P, Cappo J, Schleiermacher G, et al. Control-FREEC: a tool for assessing copy number and allelic content using next-generation sequencing data. Bioinformatics. 2012;28:423–5

66. Cortés-Ciriano I, Lee JJ-K, Xi R, Jain D, Jung YL, Yang L, et al. Comprehensive analysis of chromothripsis in 2,658 human cancers using whole-genome sequencing. Nat Genet. 2020;52:331–41

67. Sollier E, Heilmann J, Gerhauser C, Scherer M, Plass C, Lutsik P. Figeno: multi-region genomic figures with long-read support. Bioinformatics. 2024;40:btae354

68. Patel H, Ewels P, Peltzer A, Manning J, Botvinnik O, Sturm G, et al. nf-core/rnaseq: nf-core/rnaseq v3.14.0 - Hassium Honey Badger [Internet]. 2024 [cited 2024 Jul 22]. Available from: https://zenodo.org/records/10471647

69. Dobin A, Davis CA, Schlesinger F, Drenkow J, Zaleski C, Jha S, et al. STAR: ultrafast universal RNA-seq aligner. Bioinformatics. 2013;29:15–21

70. Patro R, Duggal G, Love MI, Irizarry RA, Kingsford C. Salmon provides fast and bias-aware quantification of transcript expression. Nat Methods. 2017;14:417–9

71. Love MI, Huber W, Anders S. Moderated estimation of fold change and dispersion for RNA-seq data with DESeq2. Genome Biol. 2014;15:550

72. Stephens M. False discovery rates: a new deal. Biostatistics. 2017;18:275–94

73. Ritchie ME, Phipson B, Wu D, Hu Y, Law CW, Shi W, et al. limma powers differential expression analyses for RNA-sequencing and microarray studies. Nucleic Acids Res. 2015;43:e47

74. Hao Y, Stuart T, Kowalski MH, Choudhary S, Hoffman P, Hartman A, et al. Dictionary learning for integrative, multimodal and scalable single-cell analysis. Nat Biotechnol. 2024;42:293–304

75. Butler A, Hoffman P, Smibert P, Papalexi E, Satija R. Integrating single-cell transcriptomic data across different conditions, technologies, and species. Nat Biotechnol. 2018;36:411–20

76. Kiselev VY, Yiu A, Hemberg M. scmap: projection of single-cell RNA-seq data across data sets. Nat Methods. 2018;15:359–62

77. Gao T, Soldatov R, Sarkar H, Kurkiewicz A, Biederstedt E, Loh P-R, et al. Haplotype-aware analysis of somatic copy number variations from single-cell transcriptomes. Nat Biotechnol. 2023;41:417–26

78. Dunham I, Kundaje A, Aldred SF, Collins PJ, Davis CA, Doyle F, et al. An integrated encyclopedia of DNA elements in the human genome. Nature. 2012;489:57–74

79. Zeller C, Richter D, Jurinovic V, Valtierra-Gutiérrez IA, Jayavelu AK, Mann M, et al. Adverse stem cell clones within a single patient’s tumor predict clinical outcome in AML patients. J Hematol OncolJ Hematol Oncol. 2022;15:25

80. van de Werken HJG, Landan G, Holwerda SJB, Hoichman M, Klous P, Chachik R, et al. Robust 4C-seq data analysis to screen for regulatory DNA interactions. Nat Methods. 2012;9:969–72

81. Carter B, Ku WL, Kang JY, Hu G, Perrie J, Tang Q, et al. Mapping histone modifications in low cell number and single cells using antibody-guided chromatin tagmentation (ACT-seq). Nat Commun. 2019;10:3747

82. Corces MR, Trevino AE, Hamilton EG, Greenside PG, Sinnott-Armstrong NA, Vesuna S, et al. An improved ATAC-seq protocol reduces background and enables interrogation of frozen tissues. Nat Methods. 2017;14:959–62

83. Krijger PHL, Geeven G, Bianchi V, Hilvering CRE, de Laat W. 4C-seq from beginning to end: A detailed protocol for sample preparation and data analysis. Methods. 2020;170:17–32

84. Nilsson T, Waraky A, Östlund A, Li S, Staffas A, Asp J, et al. An induced pluripotent stem cell t(7;12)(q36;p13) acute myeloid leukemia model shows high expression of MNX1 and a block in differentiation of the erythroid and megakaryocytic lineages. Int J Cancer. 2022;151:770–82

85. Wermke M, Camgoz A, Paszkowski-Rogacz M, Thieme S, von Bonin M, Dahl A, et al. RNAi profiling of primary human AML cells identifies ROCK1 as a therapeutic target and nominates fasudil as an antileukemic drug. Blood. 2015;125:3760–8

