## Supplementary Material for "Pyjacker identifies enhancer hijacking events in acute myeloid leukemia including *MNX1* activation via deletion 7q"

**Supplementary Table 1**. Methods for enhancer hijacking detection.

| Method | Required data | Can be run without matched normals | Can detect enhancer hijacking events in single samples | Uses expressi on level | Uses monoallelic expression | Uses enha ncers |
|---|---|---|---|---|---|---|
| CESAM | breakpoints + expression | YES | NO | YES | NO | NO |
| SVXpress | WGS + RNA-seq | YES | NO | YES | NO | NO |
| HYENA | WGS + RNA-seq | YES | NO | YES | NO | NO |
| cis-X | WGS + RNA-seq | NO | YES | YES | YES | YES |
| NeoLoopFi nder | HiC | YES | YES | NO | NO | NO |
| pyjacker | WGS + RNA-seq | YES | YES | YES | YES | YES |

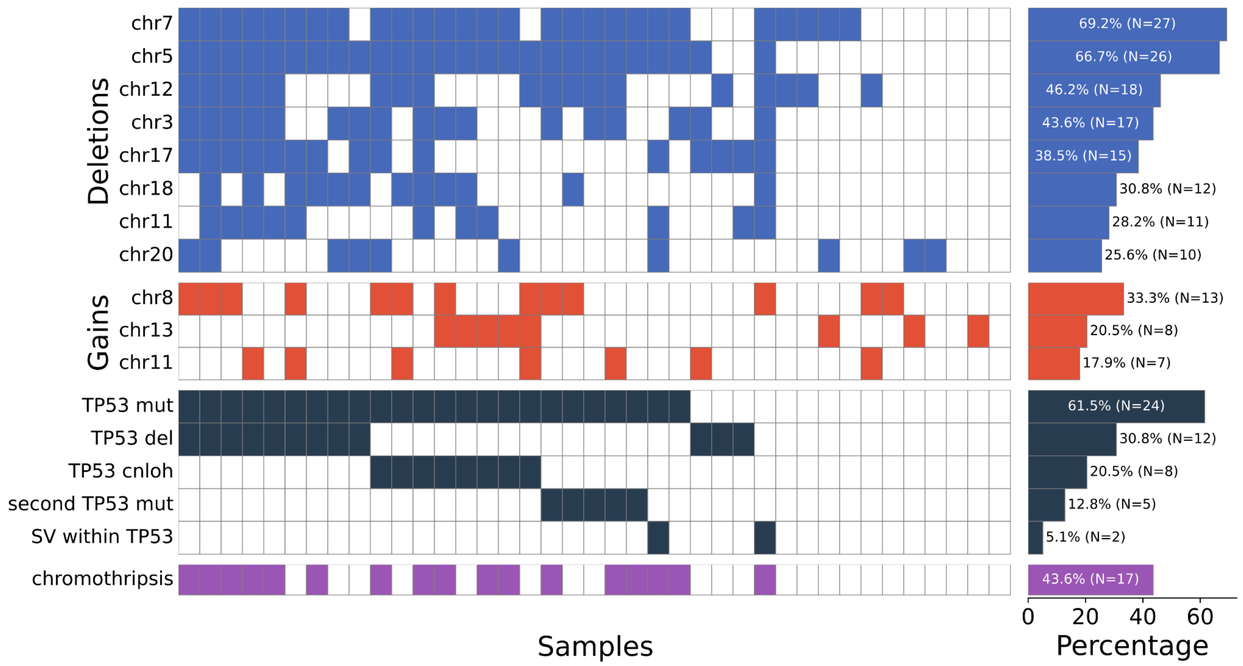


**Supplementary Figure 1. Summary of the somatic alterations in the 39 ckAML samples**. Heatmap showing the most common copy number alterations, TP53 status and chromothripsis status for the cohort of 39ckAML samples profiled with WGS and RNAseq. For copy number alterations, we counted each chromosome having at least 1Mb deleted or gained.

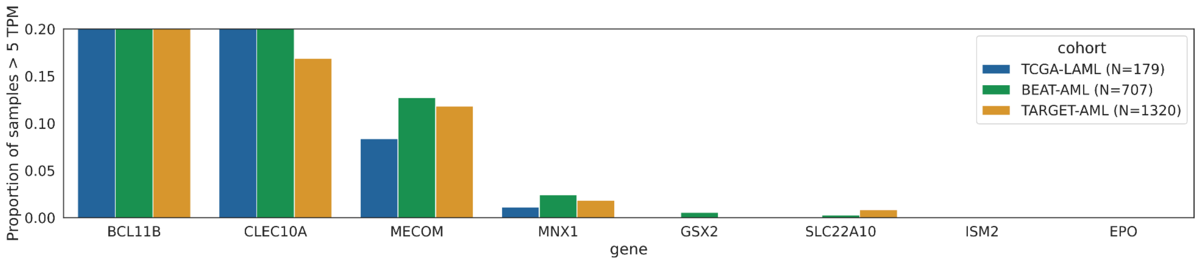


**Supplementary Figure 2. Proportion of samples expressing the top pyjacker hits, for several AML cohorts profiled with RNA-seq.** *BCL11B* and *CLEC10A* are expressed in normal T-cells and dendritic cells, respectively, which explains their expression in many samples.

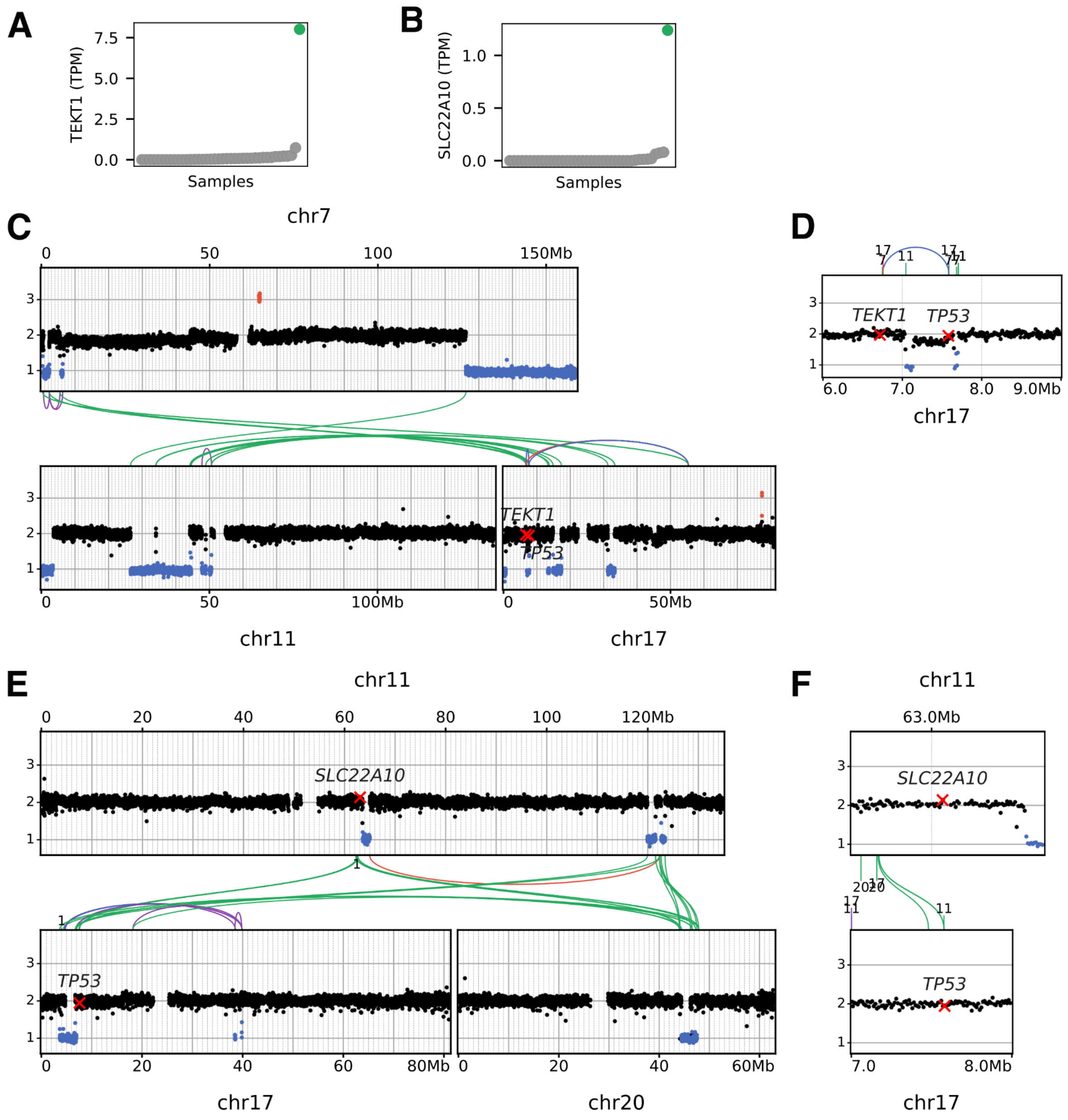


**Supplementary Figure 3. Example rearrangements leading to gene activation and TP53 inactivation. A**. *TEKT1* expression in all samples, with sample 16PB3075 (with breakpoint close to *TEKT1*) highlighted in green. **B**. *SLC22A10* expression in all samples, with sample 15KM20146 (with breakpoint close to *SLC22A10*) highlighted in green. **C**. Copy numbers and SVs on chromosomes 7,11 and 17 in sample 16PB3075. **D**. Copy numbers and SVs around *TEKT1* and *TP53* in sample 16PB3075. **E**. Copy numbers and SVs on chromosomes 11,17 and 20 in sample 15KM20146. **F**. Copy numbers and SVs around *SLC22A10* and *TP53* in sample 15KM20146.

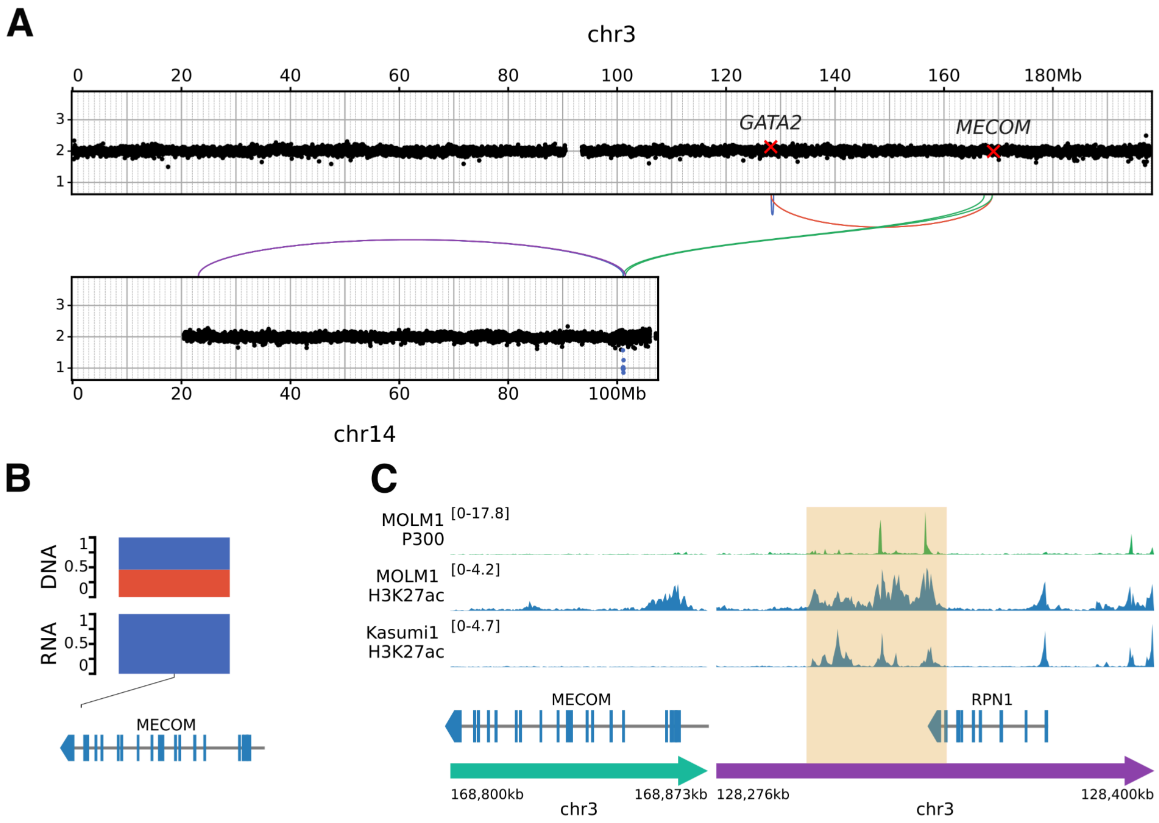


**Supplementary Figure 4. Rearrangements leading to MECOM monoallelic expression in sample 15KM20146**. **A**. Copy numbers and SVs on chr3 and chr14 for sample 15KM20146. **B**. Variant allele frequencies of SNPs in *MECOM*, in DNA and RNA of sample 15KM20146. **C**. H3K27ac and P300 tracks around the breakpoint leading to *MECOM* expression in sample 15KM20146.

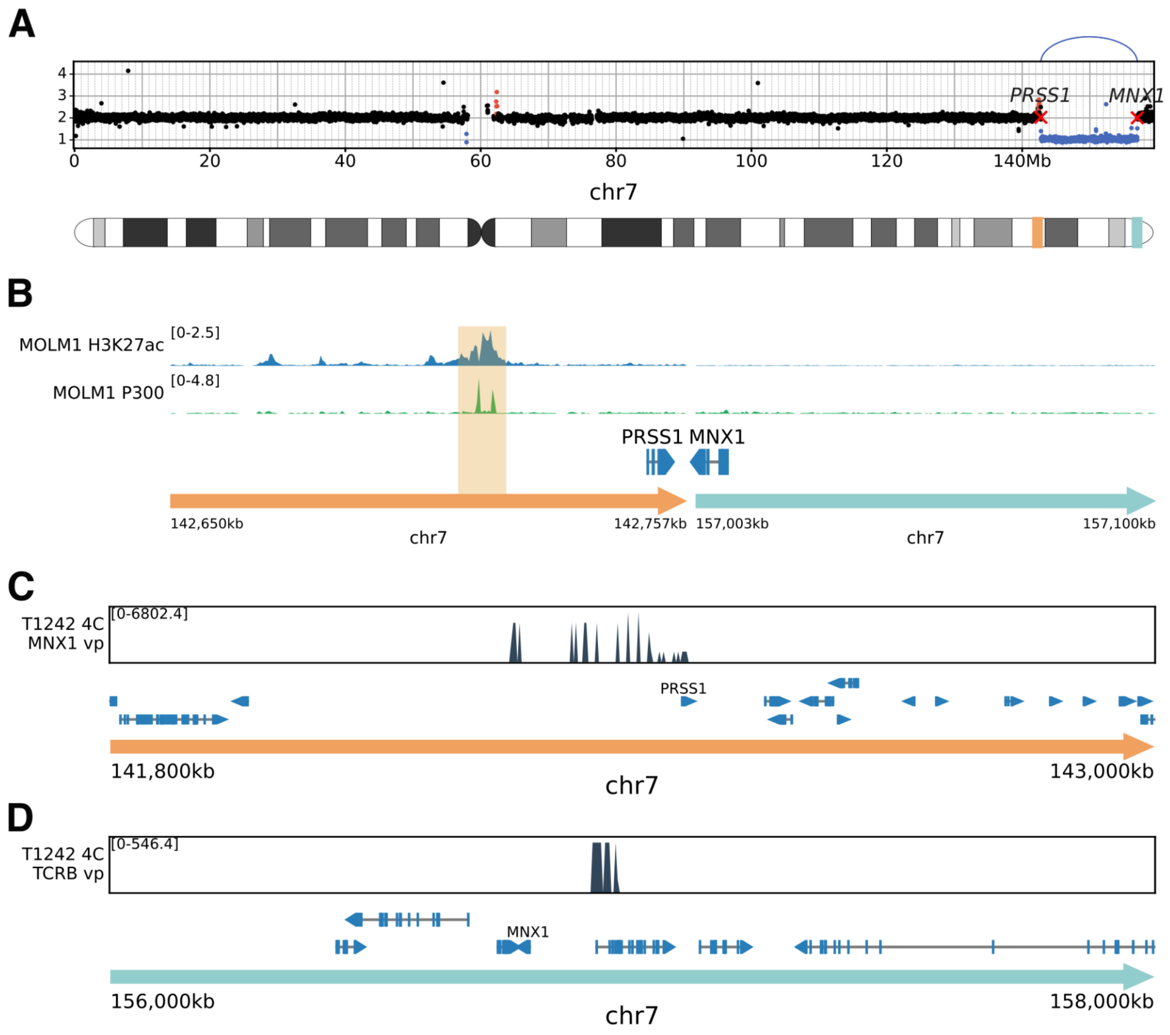


**Supplementary Figure 5. Deletion between the TCR beta locus and *MNX1* leading to *MNX1* expression. A**. Copy numbers and SVs on chr7 for sample T1242, with a smaller del(7q) leading to *MNX1* expression. Here, the coordinates are in hg38 reference because the left breakpoint (between *PRSS1* and *PRSS2*, within the T cell receptor beta locus) is in a region missing from the hg19 reference. **B**. ChIP-seq track of H3K27ac and P300 in MOLM-1 showing the putative enhancer responsible for *MNX1* activation in sample T1242 (hg38 reference). **C**. 4C track for sample T1242 with an *MNX1* viewpoint (vp), showing interaction with the TCR beta locus region (hg19 reference). **D**. 4C track for sample T1242 with a viewpoint at the TCR beta locus, showing interaction with the *MNX1* region (hg19 reference).

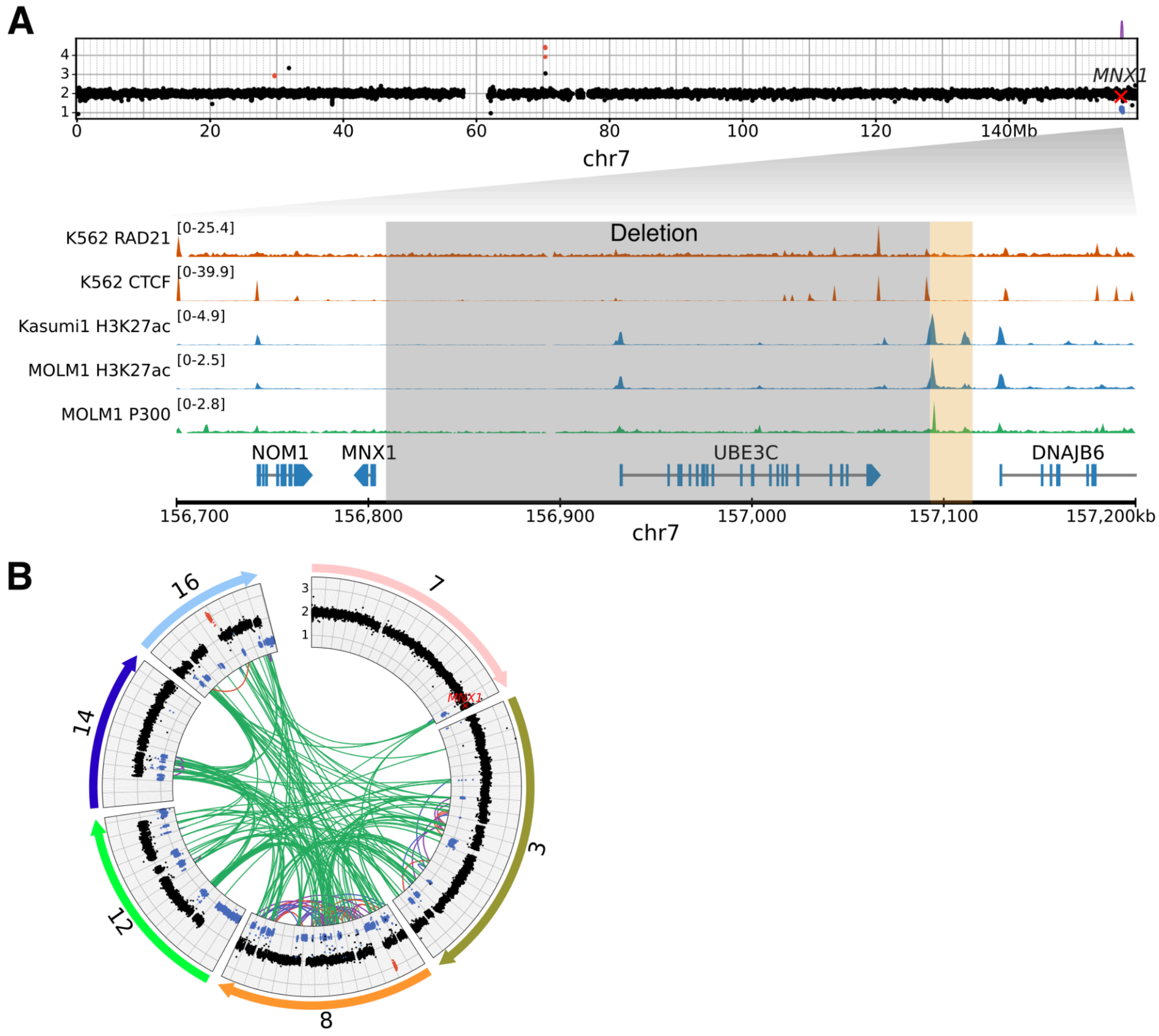


**Supplementary Figure 6. Alternative rearrangements leading to *MNX1* expression. A**. Copy numbers and SVs for sample T9058, with a 300 kb deletion to the right of *MNX1*, and ChIP-seq tracks showing the putative enhancer (highlighted in orange) to the right of the deleted region (highlighted in gray). **B**. Copy numbers and SVs in sample U4712, with a complex chromothripsis event involving multiple chromosomes, including breakpoints near *MNX1*.

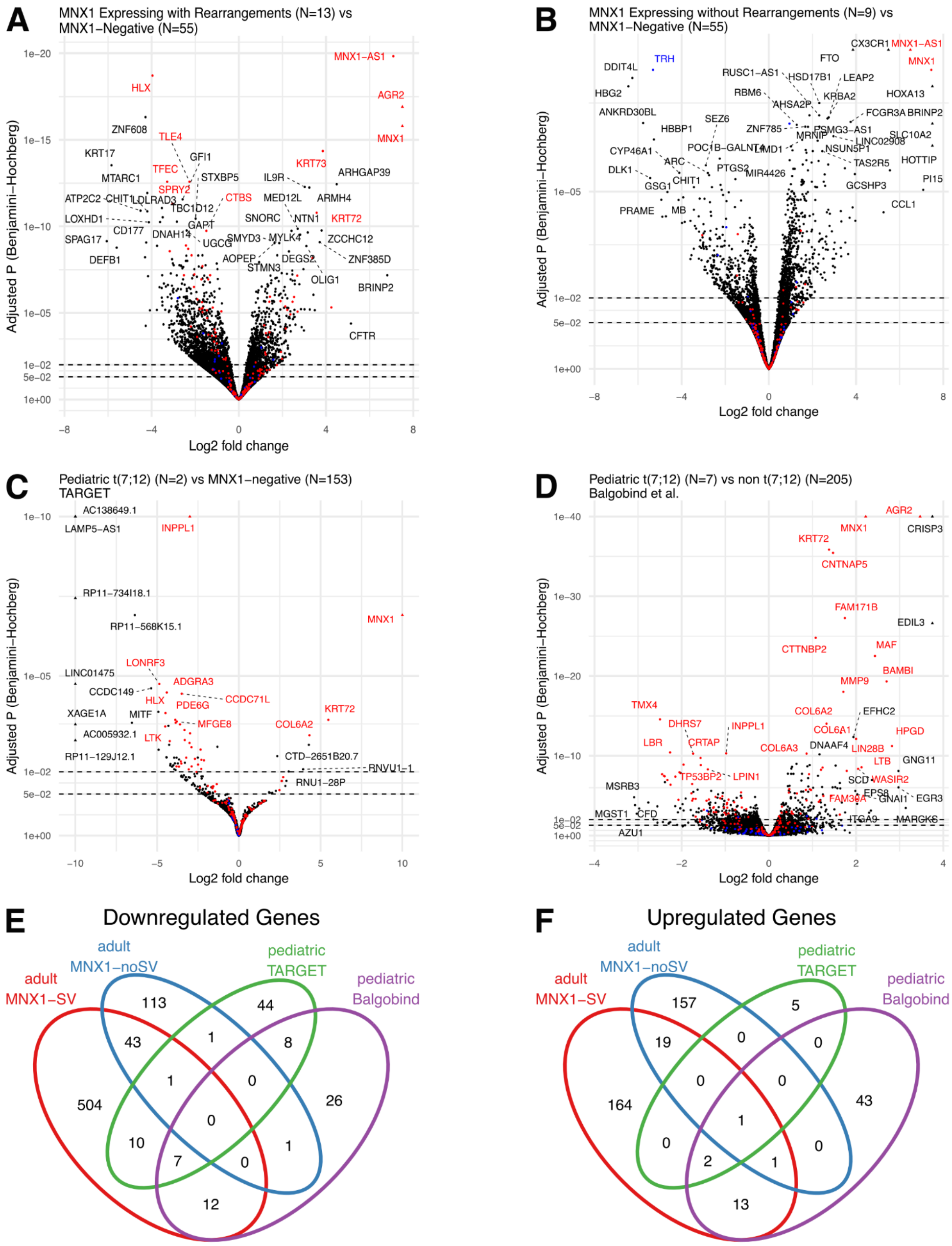


**Supplementary Figure 7. Differential gene expression analysis of *MNX1* status in adult and pediatric AML.** Genes marked in red have been proposed as an *MNX1*-associated gene signature in t(7;12) pediatric AML (PMID 36057683). Genes marked in blue have been proposed as an *NPM1*-associated gene signature (PMID 16109776). **A**. Volcano plot of adult AML with *MNX1* expression and an associated SV compared to *MNX1*-negative. **B**. Volcano plot of adult AML with *MNX1* expression without an associated SV compared to *MNX1*-negative. **C-D**. Volcano plots of pediatric AML with t(7;12)(q36;p13) compared to all other karyotypes, for two cohorts: microarray data from Balgobind et al. 2011 (**C**) and RNA-seq data from TARGET-AML (**D**). **E-F**. Intersection analysis of significantly downregulated (**E**) or upregulated (**F**) genes across the four comparisons (abs(-log10(Padj)*log(FC)>=5)).

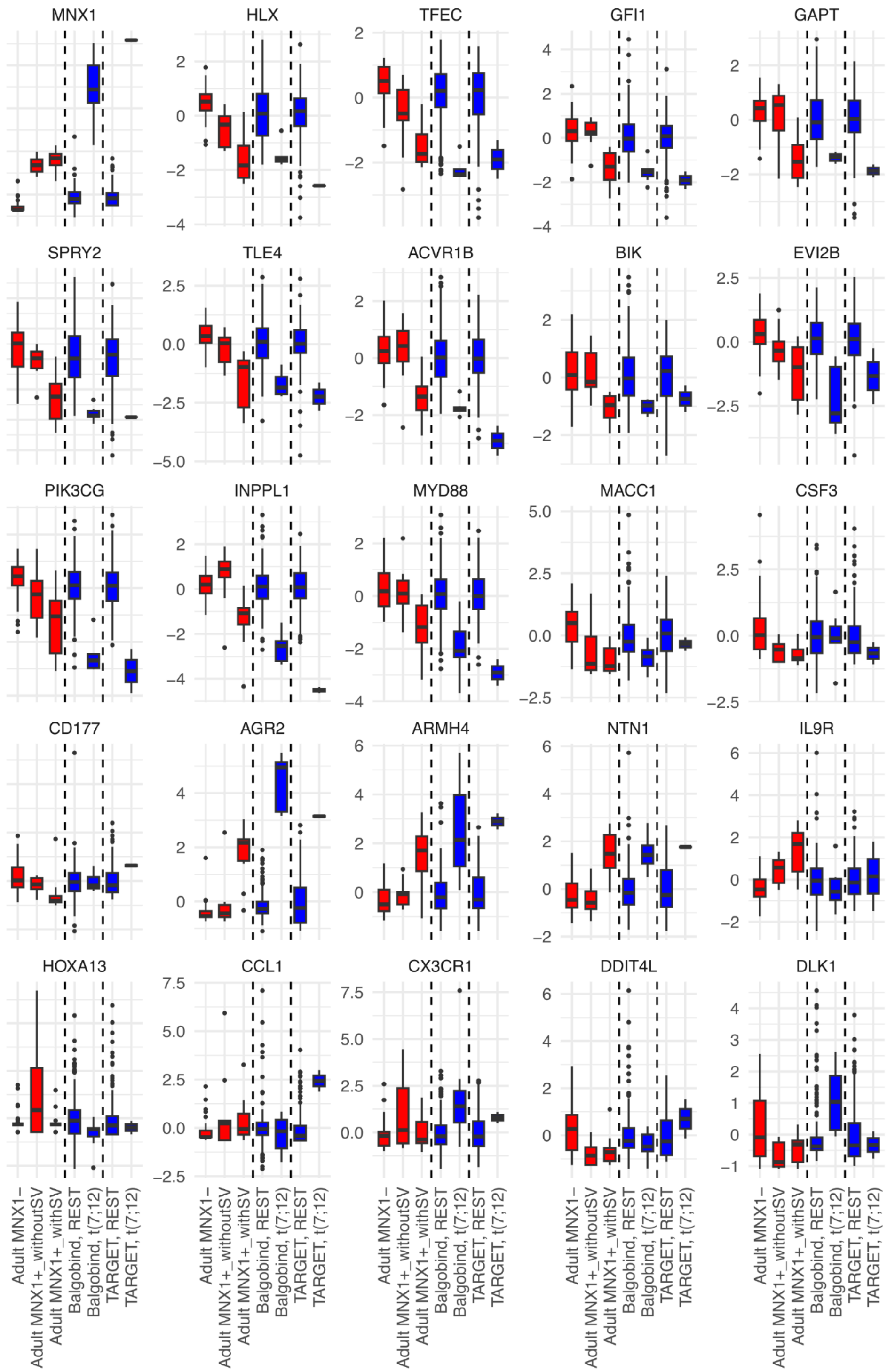


**Supplementary Figure 8. Gene expressions of 25 selected cancer and hematological development associated genes differentially expressed under *MNX1* activation.** Red boxplots represent the adult AML cohort presented in this study, whereas the blue boxplots represent the previously published Balgobind et al. and TARGET pediatric AML cohorts. The values are Z scores of vst-transformed normalized expressions.

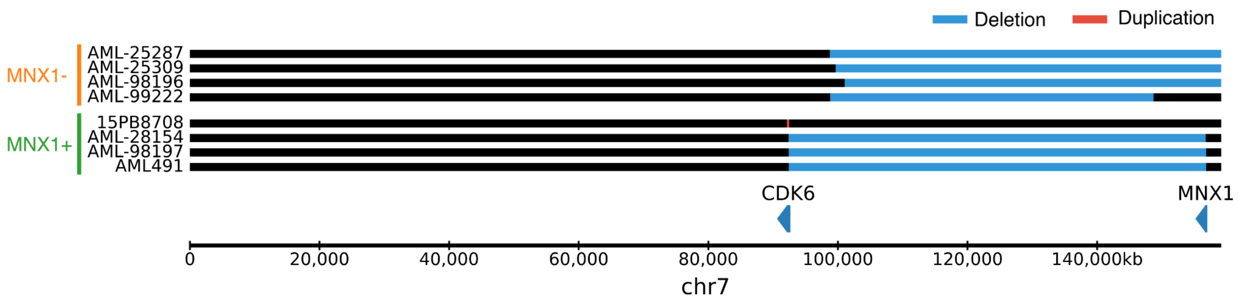


**Supplementary Figure 9**. **Copy number alterations on chromosome 7 for samples profiled with scRNA-seq.** Four *MNX1*-expressing samples (three with del(7q) and one with the *CDK6* enhancer duplicated next to *MNX1*) were profiled, as well as four control samples with del(7q) but with different breakpoints, not resulting in *MNX1* activation.

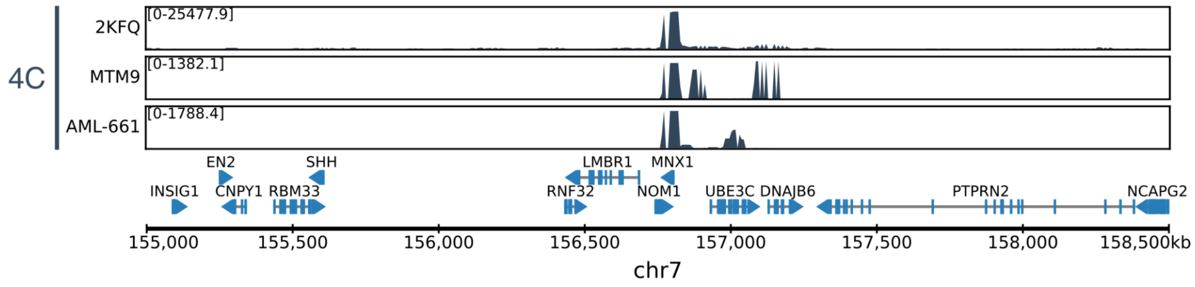


**Supplementary Figure 10**. **Reciprocal 4C.** 4C data using as viewpoint the *CDK6* region (chr7:92268000 hg19), shown in the region around *MNX1*, for del(7q) samples: MTM9 is an AML patient sample, AML-661 is a PDX sample derived from an AML patient with del(7q).

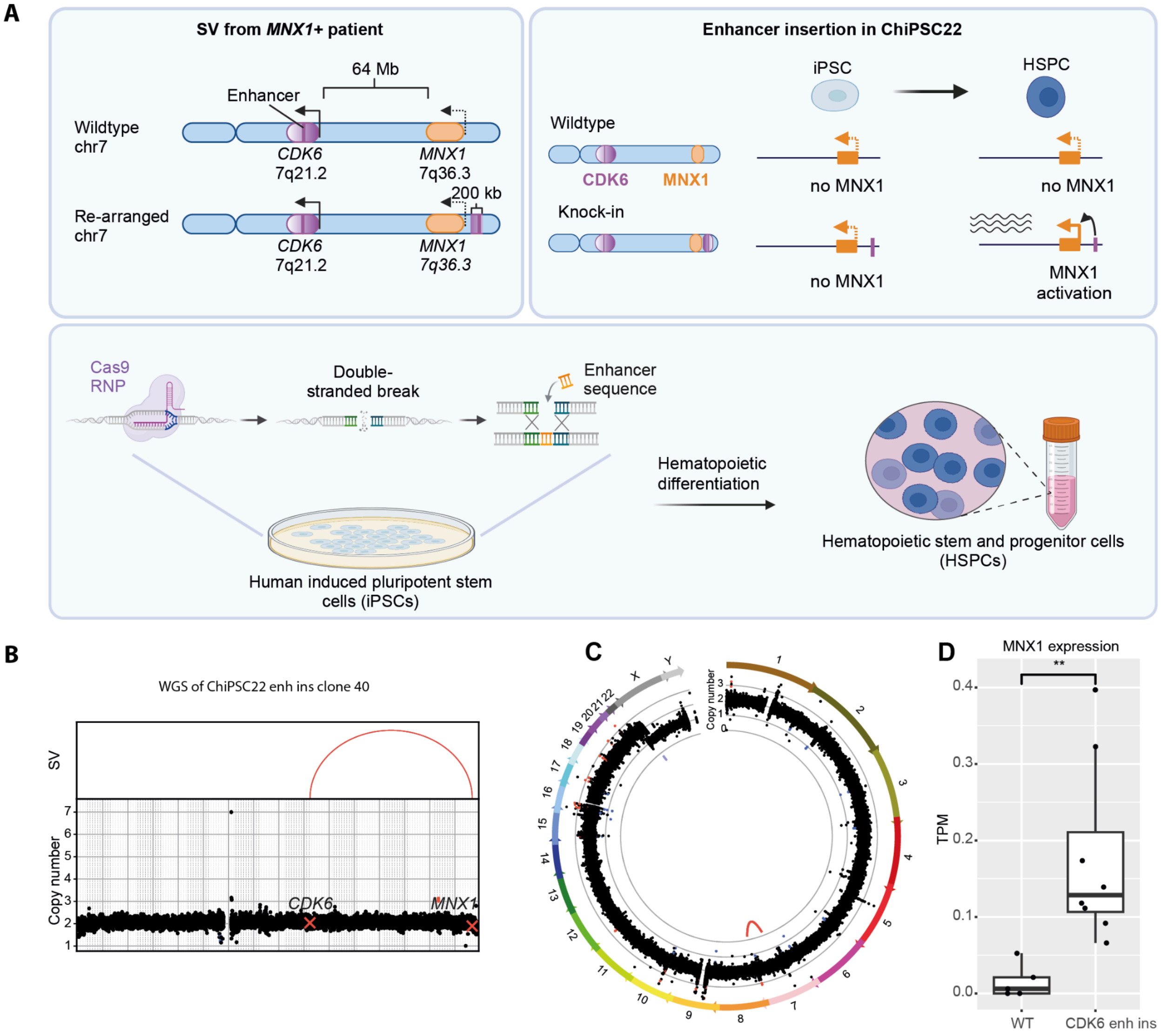


**Supplementary Figure 11. Insertion of enhancer candidate in human iPSC line. A.** Schematic overview of experimental design: A 1 kb region containing a putative enhancer was inserted upstream of the *MNX1* promoter in ChiPSC22 cells using CRISPR/Cas9. Upon differentiation of the edited iPSCs into HSPCs, the enhancer could interact with the *MNX1* promoter resulting in activated transcription. **B.** Copy number variation on chromosome 7 in the engineered cell line validating the insertion of the 1 kb region. **C.** Circos plot for the same cell line showing the absence of other rearrangements. **D.** *MNX1* expression in TPM for the parental ChiPSC22 HSPCs (n=5, from independent differentiation experiments) compared to the engineered cell with the enhancer insertion (n=8, from independent differentiation experiments for 2 different cell lines). **P < 0.01 using t-test
